# The conical shape of DIM lipids promotes *Mycobacterium tuberculosis* infection of macrophages

**DOI:** 10.1101/649541

**Authors:** Jacques Augenstreich, Evert Haanappel, Guillaume Ferré, George Czaplicki, Franck Jolibois, Nicolas Destainville, Christophe Guilhot, Alain Milon, Catherine Astarie-Dequeker, Matthieu Chavent

## Abstract

Phthiocerol dimycocerosate (DIM) is a major virulence factor of the pathogen *Mycobacterium tuberculosis* (*Mtb*). While this lipid promotes the entry of *Mtb* into macrophages, which occurs via phagocytosis, its molecular mechanism of action is unknown. Here, we combined biophysical, cell biology, and modelling approaches to reveal the molecular mechanism of DIM action on macrophage membranes leading to the first step of *Mtb* infection. MALDI-TOF mass spectrometry showed that DIM molecules are transferred from the *Mtb* envelope to macrophage membranes during infection. Multi-scale molecular modeling and ^31^P-NMR experiments revealed that DIM adopts a conical shape in membranes and aggregate in the stalks formed between two opposing lipid bilayers. Infection of macrophages pre-treated with lipids of various shapes uncovered a general role for conical lipids in promoting phagocytosis. Taken together, these results reveal how the molecular shape of a mycobacterial lipid can modulate the biological function of macrophages.

## INTRODUCTION

Phthiocerol dimycocerosate (DIM/PDIM) are highly hydrophobic lipids containing two multiple-methyl-branched fatty acid chains (**Fig. 1-a**). These lipids are mostly found in the cell wall of pathogenic mycobacteria and are particularly abundant in *Mycobacterium tuberculosis* (*Mtb*)^1^, the causative agent of tuberculosis. They constitute one of the main *Mtb* virulence factors^2^. Indeed, *Mtb* strains lacking DIM are drastically attenuated^3^ and are more likely to be killed by the early pulmonary innate immune response^4^, when the bacteria encounter macrophages. Recent work has revealed that DIM modulate macrophage metabolism^5^ and immune functions^6, 7^. In particular, DIM increase the ability of *Mtb* to infect macrophages by modulating phagocytosis^8^, a fundamental immune process involving membrane remodeling. However, how DIM intervene in these cellular processes remains poorly understood.

**Figure 1:**
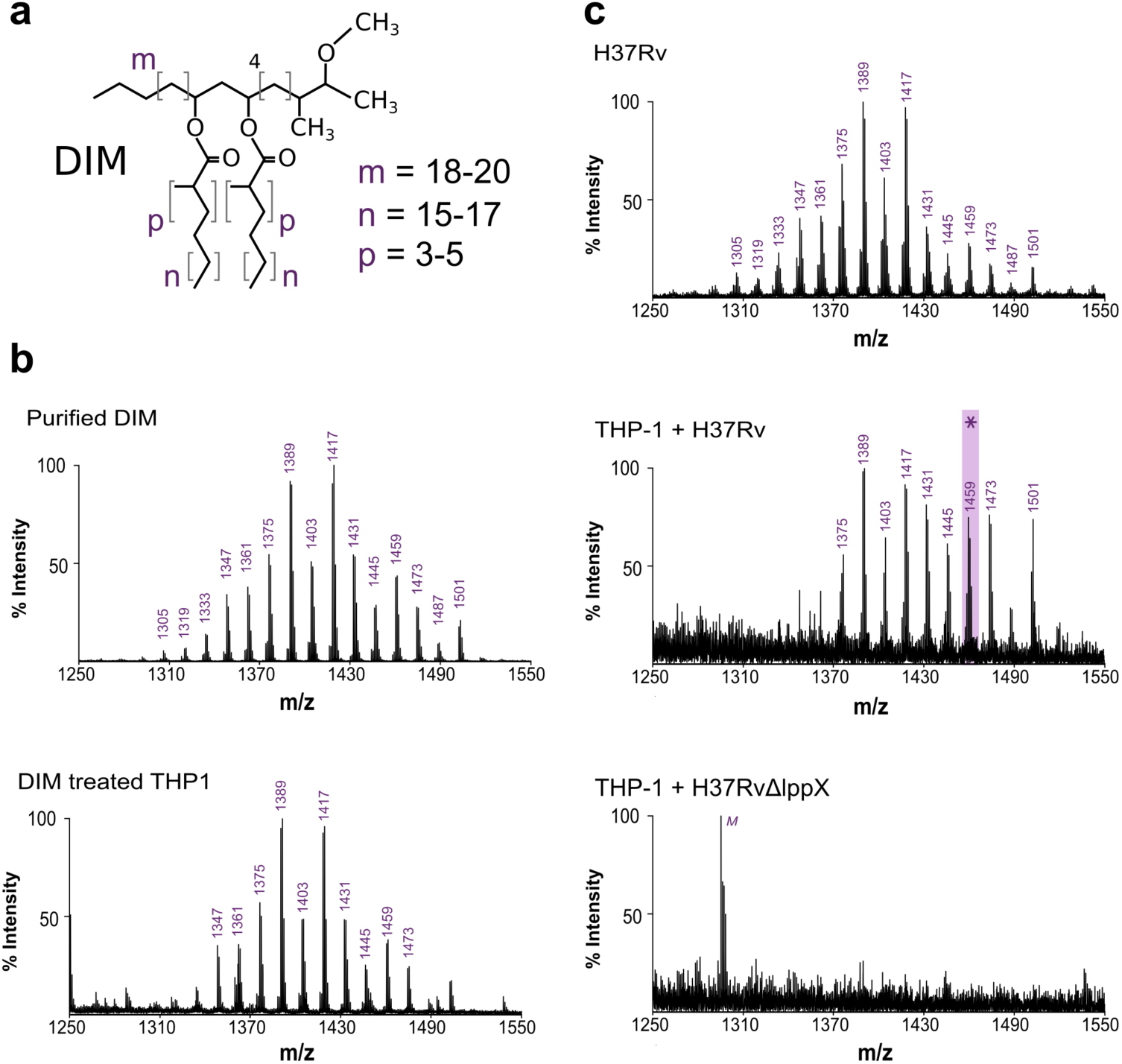
DIM are transferred from the bacterial envelope to macrophage membranes. **(a)** Structure of the DIM family of lipids, where m denotes the range of carbon atoms on the phthiocerol moiety, and n and p on the mycocerosate moieties. **(b)** MALDI-TOF mass spectra of purified DIM and of the membrane fraction of macrophages treated with DIM. **(c)** MALDI-TOF mass spectra of WT Mtb (HR37v) and of the membrane fraction of macrophages infected by H37Rv or by the H37RvΔlppX mutant. M: low intensity peak corresponding to the detection of the matrix molecule in the DIM region of interest. The star symbol highlights the mass of the DIM molecule chosen for the modelling, with m=18, n=17, and p=4.

*Mtb* synthesizes a large variety of lipid virulence factors, most of which are amphipathic glycolipids. These glycolipids act through their saccharide domains as potential ligands for membrane receptors on macrophages to induce *Mtb* phagocytosis^9^. Lacking a saccharide moiety, DIM cannot engage in such interactions. In contrast, the molecular mechanism involving DIM may be related to a global effect on the physical properties of the host cell membrane, such as its fluidity and organization^8^. Modifying such properties can be a successful strategy for bacteria to modulate eukaryotic cell functions. Several types of pathogenic mycobacteria apply this strategy to influence the fate of their host cells. For example, *M. ulcerans* produces the lipid-like endotoxin mycolactone which interacts with host membranes and disturbs their lipid organization^10^. In addition, pathogenic mycobacteria use lipoarabinomannan to enter neutrophils and prevent phagolysosome formation^11^.

The biophysical properties of DIM in biological membranes not yet been characterized at the molecular level. In particular, it is unclear if such a complex and large lipid can be incorporated in a simple phospholipid bilayer, and what shape DIM must adopt in such a membrane. The shape of lipid molecules, determined by structural properties^12^ like their head group size, acyl chain lengths and degrees of acyl chain unsaturation, can drastically affect the structure and organization of biological membranes^13, 14^. Studying how the molecular shape of lipids may disorganize lipid bilayers and how this can be related to biological function is still a challenge^15^. It requires linking the structure of molecules and their biophysical actions at the nanoscale to macroscopic consequences on the cell functions. To achieve this for DIM, we developed a multidisciplinary approach combining multiscale Molecular Dynamics (MD) simulations, solid-state NMR and cell biology experiments. This revealed how the molecular shape of DIM can affect macrophage membranes to promote phagocytosis.

## RESULTS

### DIM are transferred to host cell membranes during macrophage infection

First, we used MALDI-TOF mass spectrometry to assess whether DIM added to host cells is incorporated into their membranes. Human macrophage (THP-1) cells were treated with purified DIM, and the mass spectrum of the extracted lipids was compared with the spectrum of purified DIM. The structure of DIM consists of a long chain of phthiocerol (3-methoxy, 4-methyl, 9,11-dihydroxy glycol) esterified with two mycocerosic acids (long-chain multiple-methyl-branched fatty acids) (**Fig. 1-a**). In agreement with the MycoMass database^16^, the purified DIM mass spectrum is characterized by a cluster of pseudomolecular ions [M + Na]^+^ between m/z = 1305 and m/z = 1501, in increments of m/z = 14 (**Fig. 1-b**) reflecting the variability of chain lengths and methylations of the molecule. We observed that the spectrum of the extracted lipids from DIM-treated THP-1 cells showed a very similar series of peaks to that of purified DIM (**Fig. 1-b**). These peaks were absent in the spectrum of lipid extracts from untreated cells (**Fig. S1-a**). Hence, exogenously delivered DIM can be inserted into macrophage membranes and were detectable by MALDI-TOF mass spectrometry.

We next investigated if DIM could be transferred from the *Mtb* envelope to macrophage membranes during infection. To test this, we infected THP-1 macrophages with the WT *Mtb* strain H37Rv for 2 h at a multiplicity of infection (MOI) of 15:1. At 40 h post-infection, the membrane fraction of the infected macrophages showed a mass spectrum similar to the lipid signature of DIM isolated from the H37Rv inoculum (**Fig. 1c**). We noticed a distinct shift towards longer DIM chain lengths consistent with the reported increase in molecular mass of DIM during *Mtb* infection^17^. The residual bacterial contamination of the macrophage membrane fractions was less than 1500 cfu, well below the threshold for detection of DIM extracted directly from bacteria (between 10^5^ and 10^7^ cfu, see **Fig. S1b**). Our data therefore strongly support the model that DIM is transferred from *Mtb* to the membranes of infected macrophages.

To verify whether DIM exposure at the surface of *Mtb* is required for their transfer to macrophage membranes, we infected macrophages with a mutant strain (H37RvΔlppX) lacking LppX, a lipoprotein required for the translocation of DIM to the outer membrane of *Mtb*^18^. After infection, we did not observe the typical mass spectrum of DIM in the membrane fraction of H37RvΔlppX-infected cells (**Fig. 1c**). We verified that the WT and mutant strains produced similar amounts of DIM (**Fig. S1b**).

Taken together, these results demonstrate that DIM molecules are indeed transferred to the membranes of macrophages during infection, provided they are exposed at the surface of *M. tuberculosis*.

### DIM accommodate into a bilayer membrane by adopting a conical shape

Given their long aliphatic chains and their overall hydrophobic properties, we sought to understand how DIM might physically be accommodated in a bilayer membrane. We used a multi-scale modelling approach to gain insight into the conformation of such a complex lipid embedded in a simple phospholipid bilayer.

During macrophage infection, *Mtb* produces DIM of higher molecular weight than under non-infectious conditions^17^ (**Fig. 1-c**). We therefore modelled the structure of a DIM molecule with a molecular mass of 1459 Da (star symbol in **Fig. 1-c**), *i.e.* having chain lengths and number of methylations corresponding to m=18, n=17, and p=4 (**Fig. 1-a**). 800 ns of atomistic Molecular Dynamics (MD) simulations of a single DIM molecule in a 1-palmitoyl-2-oleoyl-*sn*-glycero-3-phosphocholine (POPC) lipid bilayer revealed that DIM is deeply embedded in the membrane and may transit between the two opposing leaflets (**Fig. 2-a**). DIM oxygen atoms preferentially remained in the proximity of the POPC ester bonds while the acyl chains stretched into the membrane hydrophobic core. The very long acyl chains (containing up to 27 carbon atoms) prevented confinement of the DIM molecule to one single leaflet. Instead, DIM seemed to be accommodated within the phospholipid bilayer by extending these chains in the inter-leaflets space (see density profile in **Fig. 2a**).

**Figure 2:**
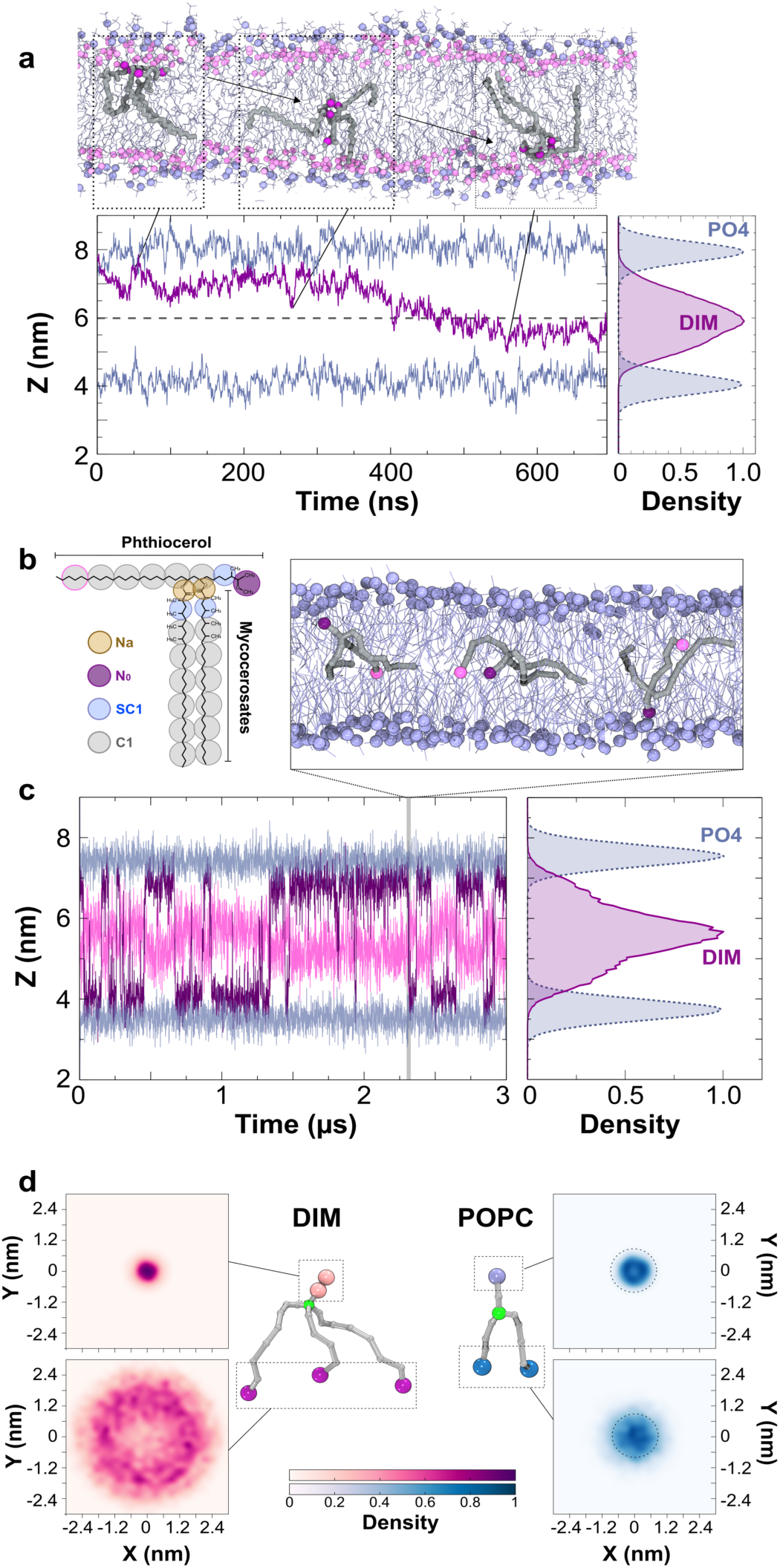
Position and shape of a single DIM molecule in a POPC bilayer. **(a)** upper inset: atomistic simulation of a DIM molecule (in gray licorice) showing its passage from one leaflet of the POPC bilayer to the opposite one. DIM oxygen atoms are represented in purple, POPC phosphorous atoms are displayed in light blue and POPC oxygen atoms in pink. Bottom left, in purple, evolution of the z-position for the center of mass of DIM’s oxygen atoms during the course of the atomistic simulation. In light blue, averaged z-position of the phosphorous atoms of the POPC molecules. Bottom right, densities of the positions of the DIM lipid and POPC phosphate groups revealing the embedded DIM position in the bilayer. **(b)** Coarse-grained model of the DIM molecule (see Methods for details). **(c)** Evolution of N0 and the last C1 particles on the phthiocerol moieties (in purple and pink, respectively) during the course of the CG simulation. This plot shows a C1 particle confined around the interleaflet space while the N0 particle stayed in the proximity of the POPC oxygen atoms. Upper inset, DIM (in grey) transit from one leaflet to the other during CG simulation. Right inset, densities of the DIM lipid and POPC phosphate groups. **(d)** 2D density projections of each extremity for DIM and POPC molecules in the x-y membrane plane (CG simulation, see also Fig. S5 displaying densities for atomistic simulations) highlighting the conical (resp. cylindrical) shape of DIM (resp. POPC) lipid. Particles depicted in green were used for molecule centering.

To further explore the dynamics of the DIM molecule within the membrane, we designed a Coarse-Grained (CG) model (**Fig. 2b**) based on the MARTINI force field (see Methods). This force field is well adapted to model a large variety of lipids and their actions on membranes and proteins^19, 20^. CG modelling of a single DIM molecule in a POPC bilayer confirmed that DIM extended its long acyl chains in between the two leaflets as seen in the atomistic simulation (**Fig. 2b, c**). Using CG modelling we were able to extend the simulation to longer time scales to see multiple DIM translocations from one leaflet to the other (**Fig. 2c**). We then increased the number of DIM molecules up to a molar DIM-to-POPC ratio of about 7% (**Fig. S4**). At low concentrations (1%, 2%, and ∼4%), DIM molecules diffused freely inside the bilayer, while at 7% they started to strongly interact with each other and form aggregates in between the two leaflets. This behavior is also observed, both experimentally and computationally, for molecules with similar structural features, like triglycerides^21, 22^.

We next sought to understand how the position of the DIM acyl chains in the inter-leaflet space affected its overall shape. To do so, we projected the positions of the lipid extremities onto the 2D membrane plane (**Fig. 2d**). When centering the molecule on the junction of the chains, this revealed very large movements of the three acyl chain extremities, while the most polar end of the phthiocerol chain remained largely static. For POPC, a similar projection displayed a completely different behavior, with comparable densities for both the headgroup and the hydrophobic acyl chain extremities (**Fig. 2d**). Comparable results were obtained from atomistic simulations (**Fig. S5**). These results can be related to the effective shape of each molecule: while it is known that POPC has a cylindrical shape, consistent with our simulations, which is suitable to form planar lipid membranes, our results indicated that DIM molecules adopt a strongly conical shape in a lipid bilayer.

### DIM drive the formation of non-bilayer membrane structures

This conical molecular shape of DIM may have important consequences for the organization of DIM-containing membranes. Indeed, conical lipids are known to destabilize the lamellar membrane phase (L*_α_*) and favor the appearance of a non-bilayer inverted-hexagonal phase (H_II_)^23^. The transition from an L*_α_*-phase to an H_II_– phase can be studied using ^31^P-NMR spectroscopy by monitoring the NMR spectra at increasing temperature (see Methods). We have employed Magic Angle Spinning (MAS) NMR spectroscopy for enhanced sensitivity. A mixture of the phospholipids 1,2-dioleoyl-*sn*-glycero-3-phosphoethanolamine (DOPE) and 1-stearoyl-2-oleoyl-*sn*-glycero-3-phosphocholine (SOPC)^24^ has been used for a range of lipids to study their propensity to induce non-bilayer phases^25, 26^. To study the influence of DIM, we used lipid membranes made of a 3:1 (mol/mol) mixture of DOPE and SOPC (see Method). With this lipid composition, the membranes remained in the L*α* phase for temperatures up to 322 K (**Fig. 3a,c**). However, incorporating 5% of DIM into the lipid mixture destabilized the L*_α_* phase, and induced a transition from a L*_α_* phase at low temperature (284 K, **Fig. 3a**) to an H_II_ phase configuration at high temperature (310 K and 322 K, **Fig. 3a**) as evidenced by the ^31^P NMR spectra. Thus, DIM destabilize the L*_α_* phase in our model membranes and promote the transition to the H_II_ phase.

**Figure 3:**
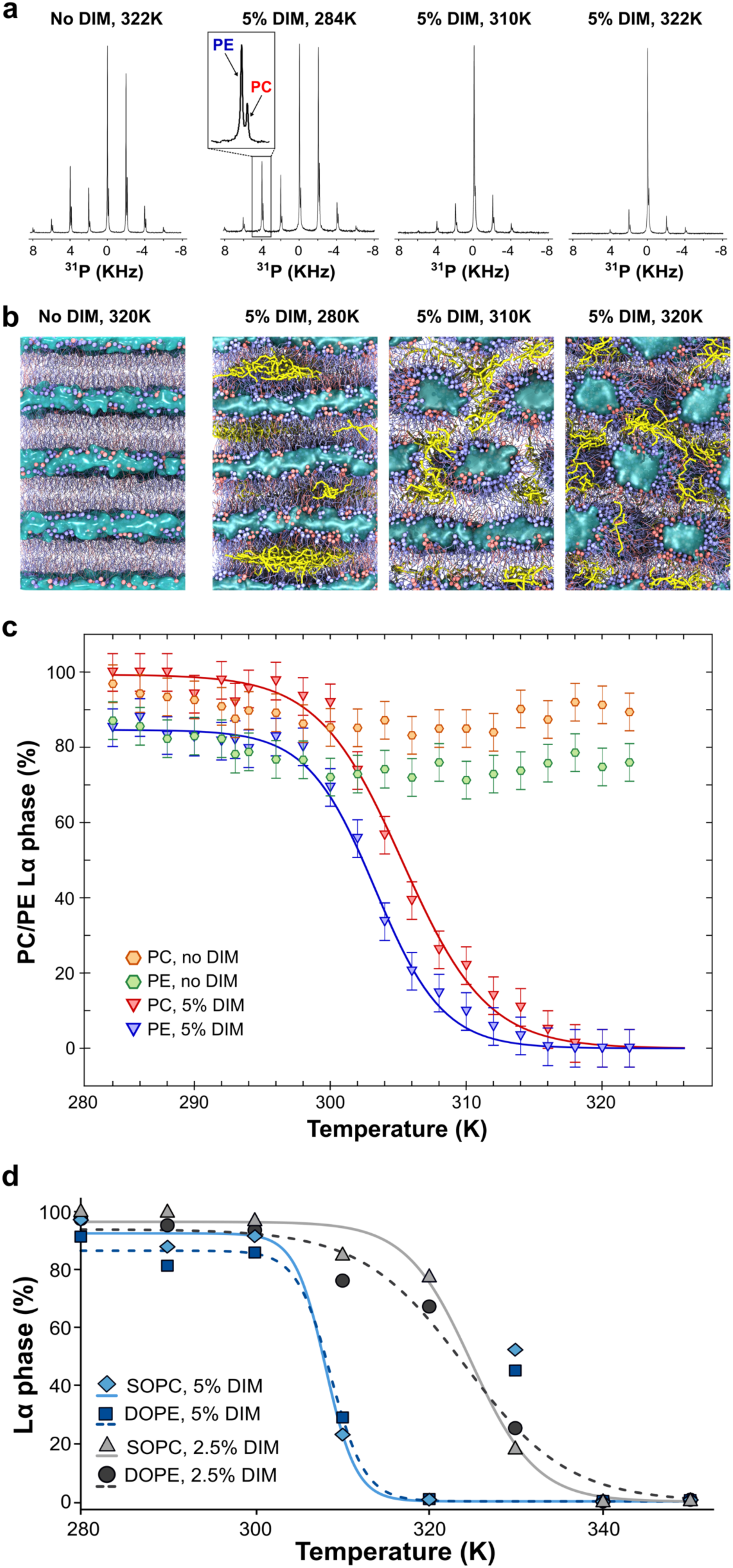
DIM induce HII phases. **(a)** 31P NMR spectra at different temperatures for DOPE/SOPC (3:1, mol/mol) without DIM or containing 5% of DIM. In the MAS ^31^P NMR spectra, each spinning sideband consists of a DOPE and a SOPC peak. **(b)** Coarse grained models of the phase transition in DOPE/SOPC (3:1, mol/mol) without DIM or containing 5% of DIM. Increasing the temperature leads to the formation of tubular structures for the DIM-containing systems while the system without DIM stay fully lamellar. Snapshots shown are taken at the end of the 3 μs simulations. SOPC molecules colored in red. DOPE molecules colored in blue. DIM molecules colored in yellow. Water molecules represented as a blue surface. **(c)** Spectral deconvolution of the ^31^P NMR spectra giving the percentage of the Lα phase as a function of the temperature for DOPE/SOPC (3:1) without DIM (orange and green hexagons) and with addition of 5% DIM (red and blue triangles). In the case of 5% DIM, the phase transition can be approximated by a sigmoid (red and blue lines). **(d)** SOPC and DOPE phase transitions calculated from CG-MD simulations for 2.5% and 5% of DIM. For the DOPE /SOPC mixture with 5% of DIM, we removed the outlier values at 330K from the curve fitting based on statistical tests (see supplementary material).

As a complementary approach to monitor the ability of DIM to induce non-bilayer phases, we performed CG-MD simulations^27^. We modelled a stack of four lipid bilayers of identical composition (3:1 DOPE/SOPC) as in the ^31^P-NMR experiments, for temperatures ranging from 280 K to 350 K (see Methods for details). Similar to the ^31^P-NMR experiments, the membranes remained in the lamellar phase in the absence of DIM (**Fig. S6**), but a temperature-driven transition occurred when 5% of DIM were added (**Fig. 3b**). These simulations allow a deeper understanding of the molecular process of the H_II_ phase transition in the presence of DIM. At low temperatures, molecules of DIM formed aggregates in the inter-leaflet space (**Fig. 3b**, 280K), as also seen in the POPC bilayer (**Fig. S4**). Increasing the temperature led to the formation of fusion stalks, hourglass-shaped lipid structures formed between neighboring bilayers. DIM aggregated in these stalks to extend their large hydrophobic tails (**Movie S1**). This aggregation stabilized and helped to increase the width of the stalks, eventually leading to the formation of tubular water-filled membrane structures (**Fig. 3b**, 310 K). At still higher temperatures (320 K and higher), DIM molecules diffused freely in the hydrophobic membrane core, thus stabilizing the H_II_ configuration (**Fig. 3b**, 320 K and **Movie S1**).

We deconvoluted the ^31^P-NMR spectra recorded between 282 K and 324 K in 2 K steps for DOPE/SOPC (3:1) with 5% of DIM, using a set of parameters obtained from reference datasets (see Methods and **Table S2)**. The ^31^P chemical shifts of DOPE and SOPC are different. Therefore, each spinning sideband in our spectra consisted of two resolved peaks (**Fig. 3a**) enabling us, in a single spectrum, to independently analyze the percentages of the L*_α_* and H_II_ phases for DOPE and for SOPC. For both lipids, we observed a continuous transition from the L*_α_* to the H_II_ phase described by a sigmoid (**Fig. 3c**, Supplementary Material and **Table S3**). The phase transition midpoint temperatures (T_50_) were 303.5 K (± 0.4) for DOPE and 305.3 K (± 0.5) for SOPC. Thus, the DOPE lipids were more susceptible to transition to an H_II_ phase than the SOPC lipids, a tendency observed in most of the tested conditions (**Table S3**). This result is in agreement with previous studies showing that conical DOPE prefers the H_II_ phase, in contrast to cylindrical POPC lipids^28^. In the CG-MD simulations, we evaluated the percentage of L*_α_* phase as a function of temperature from the distribution of lipid tilt angles (see Supplementary material), for SOPC and DOPE (**Fig. 3d**). Here, too, we observed continuous phase transitions that were fitted by a sigmoid with a midpoint transition temperature of ∼308 K (**Table S3**), which is in broad agreement with the ^31^P-NMR experiments.

CG-MD simulations revealed that the first stage of the mechanism through which DIM drive the L*_α_*-to-H_II_ phase transition involves their aggregation in membrane stalks. Hence, blocking the formation of stalks would be expected to reduce the effect of DIM. To test this, and validate our hypothesis, we replaced a fraction of SOPC with lysophosphatidylcholine (lysoPC), a lipid known to hinder the formation of fusion stalks^29, 30^. ^31^P-NMR experiments on liposomes of DOPE/SOPC/lysoPC (75:25:0), (75:20:5), and (75:15:10), each containing 2.5% of DIM, revealed an increase of the transition midpoint temperature T_50_ (**Fig. S8**) with increasing percentage of lysoPC, highlighting a diminished effect of DIM when fusion stalk formation is inhibited. We next performed CG-MD simulations to understand the molecular process involved. Consistent with the NMR experiment, including lysoPC in the simulation increased the value of T_50_ (**Fig. S8a**). From these simulations, we observed that the effect of lysoPC did not involve a direct interaction with DIM, as lysoPC was spread throughout the membranes (**Fig. S8b**). Indeed, ^31^P-NMR experiments on DOPE/SOPC (5:1, mol/mol), which displays an L*_α_*-to-H_II_ transition without DIM, also showed an increase in T_50_ upon replacing a fraction of SOPC by lysoPC (**Fig. S7**). Thus, limiting the formation of stalks decreased the ability of DIM to effectively drive the formation of non-bilayer membrane structures.

Altogether, the combination of ^31^P-NMR and CG-MD simulations revealed the ability of DIM lipids to perturb membrane organization by promoting a phase transition from the lamellar to the inverted hexagonal phase. The molecular mechanism involves an initial aggregation of DIM lipids in fusion stalks, which then leads to a complete destabilization of the lamellar phase in favor of the inverted hexagonal phase.

### High potency of DIM to induce non-bilayer phase in comparison to other lipids

We compared the ability (potency) of DIM to induce the H_II_ phase to that of lipids with different structural features using ^31^P-NMR. We first tested the effect of the concentration of DIM on the formation of the H_II_ phase. **Figure 4-b** shows that decreasing the DIM concentration to 2.5% and 1% still led to the formation of the full H_II_ phase, albeit at a higher temperature. The increased transition midpoint temperature (**Fig. 4b**) reveals a dose-response relationship, which is also observed in CG-MD simulations **Fig. 3d**). We then tested the effect of the triglyceride tripalmitin (**Fig. 4a**), which, like DIM, has three acyl chains. However, incorporation of either 2.5% or 5% of tripalmitin did not induce a full H_II_ phase transition (**Fig. 4b**). Thus, the effect of DIM on non-bilayer structure formation did not seem to be uniquely related to its three-legged structure.

**Figure 4:**
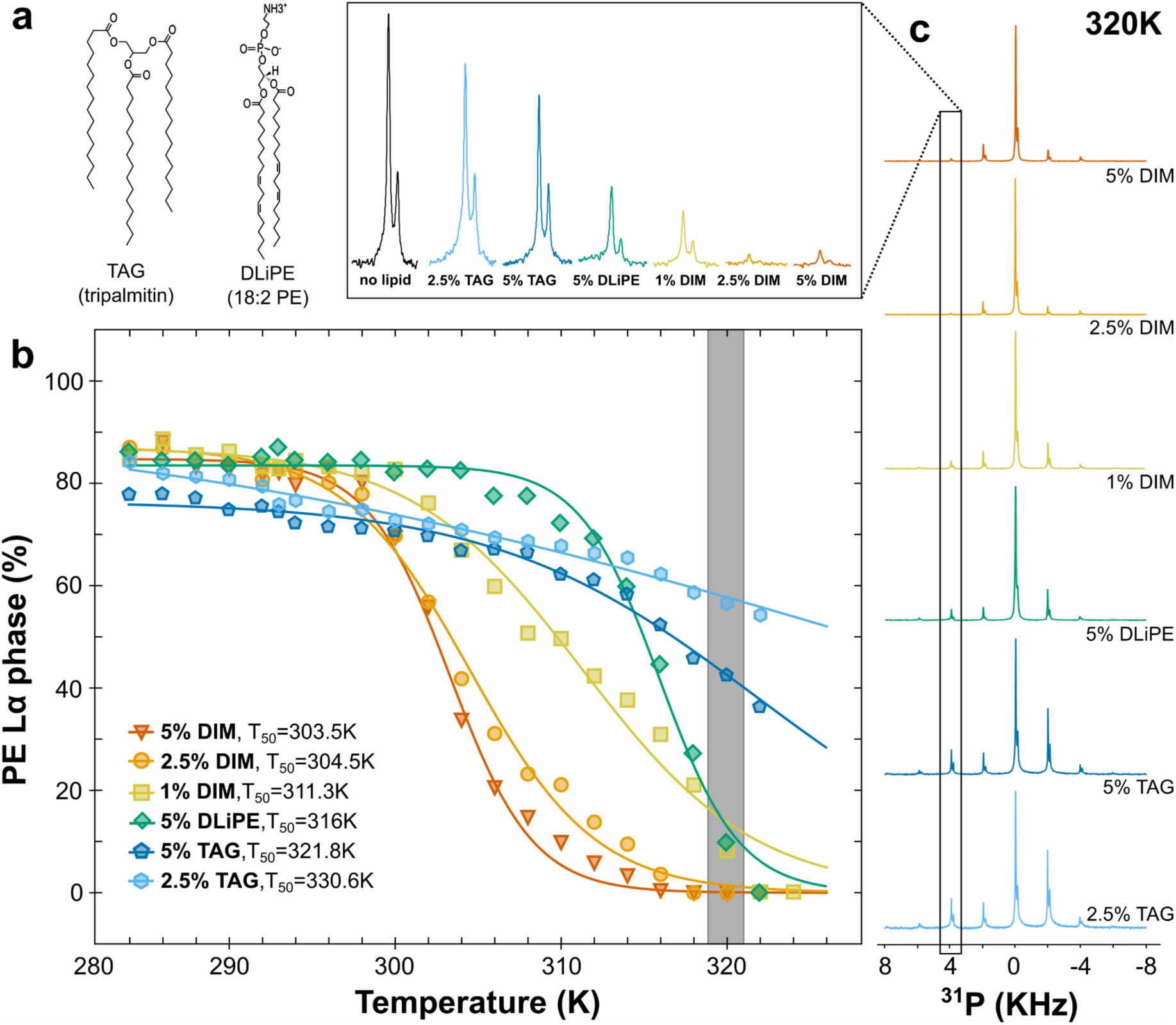
Comparison of DIM potency to induce non-bilayer phase with lipids of different shapes. **(a)** Molecular structures of the triacylglycerol (TAG) tripalmitin and of 1,2-dilinoleoyl-sn-glycero-3-phosphoethanolamine (DLiPE). **(b)** Evolution of the Lα-to-H_II_ phase transition for the DOPE molecules as a function of temperature for DOPE/SOPC (3:1) containing different concentrations of DIM, DLiPE and TAG (see also Fig. S9 for the respective curves for SOPC molecules). For clarity, error bars are omitted. As seen in Fig. 3, the error was evaluated to ±5%. The gray bar represents the points obtained from the spectra highlighted in c. (c) ^31^P NMR spectra for the lipid mixtures containing TAG, DLiPE or DIM at 320 K. The second rotation band (4 kHz), which is strongly related to the evolution of the Lα phase, is magnified in the upper panel. For comparison, the black peak depicts the second rotation band of DOPE/SOPC (3:1) at 320 K.

We finally analyzed the ability of 1,2-dilinoleoyl-*sn*-glycero-3-phosphoethanolamine (DLiPE) to induce the L*_α_*-to-H_II_ phase transition and compared it with DIM. With two double bonds in each acyl chain (**Fig. 4a**), DLiPE is expected to be a strong enhancer of hexagonal phase formation^31^. With 5% of DLiPE, we observed a full H_II_ phase transition already at 322K (**Fig. 4b,c**) and a transition midpoint temperature for DOPE of 316 K (± 0.4). However, this value is higher than for 5% DIM (303.5K ± 0.4), 2.5% DIM (304.5K ± 0.5) and even 1% DIM (311.3K ± 1.0) (**Table S3**). Thus, DIM molecules are strong inducers of non-bilayer phases, even at low concentrations.

Lipid shape can be assessed by studying inverted hexagonal phase in different lipid mixtures^32^. Here, using the L*_α_*-to-H_II_ phase transition temperature as a measure to assess lipid conical shape, we ranked the shape of the different molecules: DIM were strongly conical, DLiPE were less conical, while the tripalmitin were the least conical.

### Lipid shape modulates the entry of *Mtb* and zymosan particles into macrophages

In our previous experiments, the DIM-deficient mutant H37Rv*ΔppsE* appeared to infect macrophages with a lower efficiency than the WT strain. Coating these DIM-deficient mutant bacteria with DIM restored the WT phenotype while coating mutants with tripalmitin did not have an effect^8^. These results may now be related to their respective conical shapes: strongly conical for DIM and less conical for tripalmitin. To test whether it is specifically the conical shape of DIM that helps *Mtb* to invade macrophages, we evaluated the impact of exogenously added DIM and various other lipids on the capacity of this mutant to invade macrophages in comparison to the WT H37Rv strain (**Fig. 5a**). We confirmed that the DIM-deficient mutant infected a lower percentage of macrophages than the WT strain (**Fig. 5b,c**). Pre-treatment of macrophages with DIM restored the percentage of infected cells to a level comparable to that observed with the WT H37Rv strain in untreated macrophages (**Fig. 5b,c**). Notably, treating macrophages with the conical phospholipid POPE also enhanced the percentage of macrophages infected with the H37Rv*ΔppsE* mutant, whereas treatment with the cylindrical lipid POPC had no significant effect (**Fig. 5c**). These data support the hypothesis that the conical shape of DIM, which induces non-bilayer membrane structures, increases the efficiency of *Mtb* to infect macrophages.

**Figure 5:**
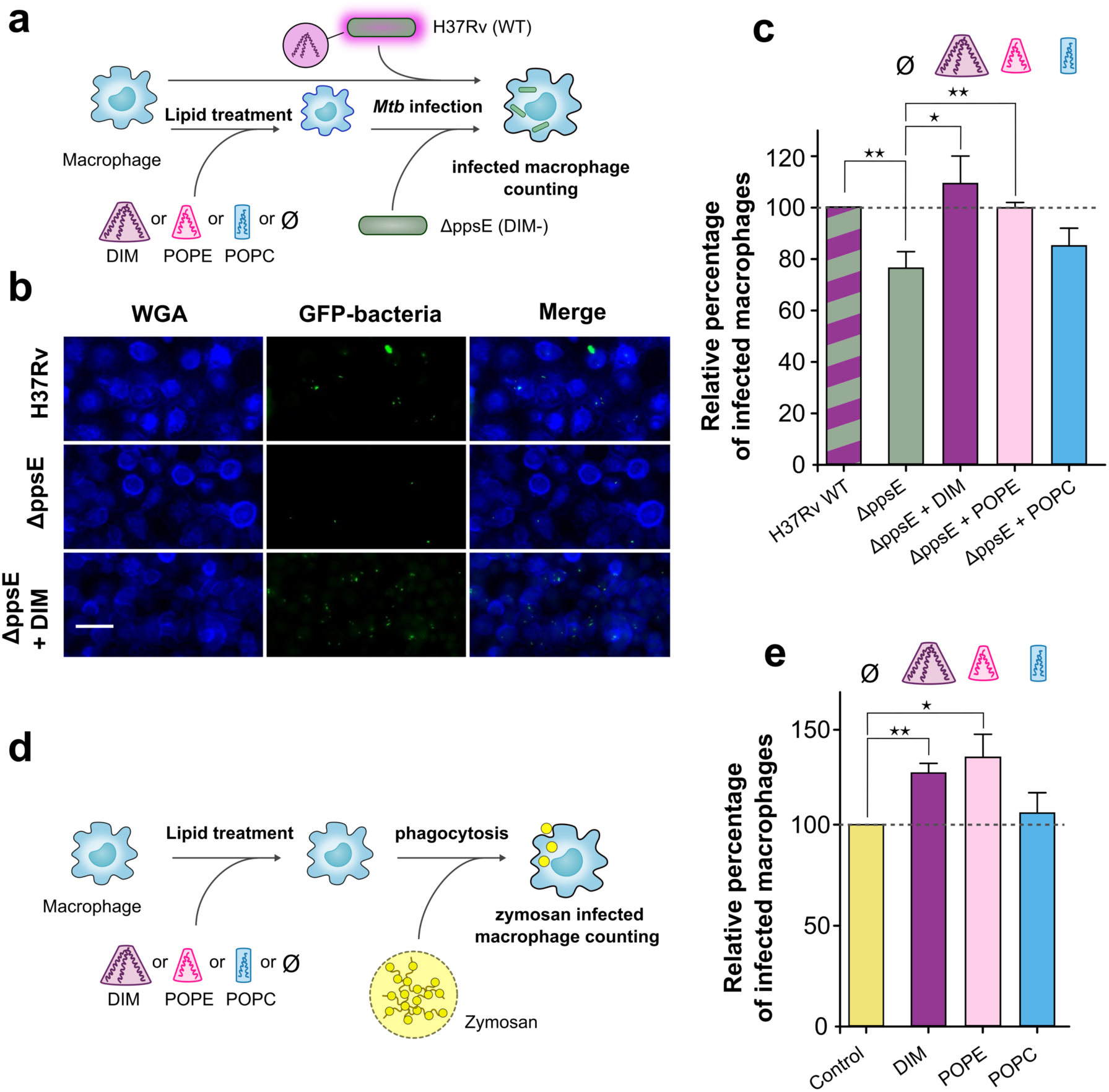
Lipid shapes modulate the entry of the MtbΔppsE mutant and zymosan into macrophages. (a) Macrophages were incubated at 37°C for 1 h with lipid solvent (Ø) or 70 μM lipids (DIM, POPC and POPE) and subsequently exposed to GFP-expressing H37Rv WT or ΔppsE (MOI 10:1) for 1 h. Cells were then fixed and processed for the quantification of infected macrophages by fluorescence microscopy. (b) Representative fluorescence microscopy images of untreated or DIM-treated macrophages stained with WGA (membrane marker, blue) and infected with H37Rv WT or ΔppsE (green); scale bar: 30 μm. (c) The histogram represents the percentage of macrophages infected with ΔppsE in lipid-treated and untreated macrophages, expressed with respect to H37Rv WT (100%). (d) Macrophages were incubated with solvent (Ø) or lipids, and then put in contact for 1 h with zymosan particles (MOI 30:1). (e) Percentages of macrophages infected with zymosan in untreated cells or cells treated with lipids. Values are expressed with respect to the uptake of zymosan in untreated cells (100%). The values are means + SEM of 8-10 separate experiments. The significance of difference in the percentage of macrophage infection between H37Rv WT and ΔppsE or between untreated and lipid-treated cells were evaluated, *, p <0.05; **, p<0.015.

To determine whether this effect of DIM on macrophages is restricted to infection by *Mtb*, we examined the effect of DIM and other lipids on the uptake of zymosan (**Fig. 5d**) a fungal polysaccharide frequently used to study non-opsonic phagocytosis^32^. We found that macrophage pre-incubation with DIM also increased zymosan uptake by macrophages in comparison to untreated conditions, as did pre-incubation with POPE but not with POPC (**Fig. 5e**). These data indicate that DIM and other conical lipids generally promote phagocytosis by macrophages.

## DISCUSSION

While lipid transfer from *Mtb* to the macrophage membranes during infection had been demonstrated for glycolipids^33^, this was never shown for DIM. By using MALDI-TOF mass spectrometry, we established that DIM molecules exposed at the envelope of *Mtb* are indeed transferred to the macrophage membranes during infection (**Fig. 1**). We envision two mechanisms that could account for this process. DIM may be exchanged by direct contact between the surface of the bacteria and the macrophage membrane at contact sites. Such direct exchange of cholesterol and cholesterol-glycolipids has been observed between *Borrelia burgdorferi* and HeLa cells^34^. Alternatively, DIM could be transported in the membranes of extracellular vesicles, which are known to be excreted by *Mtb* and other mycobacteria^35^, followed by fusion of these vesicles with the plasma membrane of the macrophages. Lipid exchange mediated by vesicle fusion was shown for *Borrelia burgdorferi*^34^ and for *Pseudomonas aeruginosa*^36^. For the latter process, lipid transfer could be favored by the conical shape, which promotes the fusion of vesicles with the host cell membrane^37^. Here, by combining ^31^P-NMR with MD simulations, we demonstrated that DIM can adopt such a conical shape and promote the formation of non-bilayer (inverted hexagonal) membrane phases (**Fig. 3**), structures important for efficient membrane fusion^38^. Membrane fusion may also be important for sealing of the phagosomal membrane during the ultimate stage of phagocytosis. Notably, our results showed that even DLiPE, a strong enhancer of non-bilayer phases, did not match the strength of DIM in promoting non-bilayer structures (**Fig. 4**). This may be explained by the fact that DIM lipids preferentially aggregate in transient stalks (**Fig. 3b and movie S1**) to stabilize them, thereby enhancing non-bilayer phase formation. Based on our modelling, the conical shape of DIM can be related to the accommodation of the very long acyl-chains of DIM to phospholipid bilayers (**Fig. 2**). This shape may also be adopted by other long acyl chain lipids that are important for *Mtb* infection^2^ and are transferred to the host cell membrane^33^ such as trehalose mono- and di-mycolate and the phenolic glycolipids, molecules structurally related to DIM.

There is now ample evidence that lipids can modulate membrane protein function^39, 40^. As seen for lipids such as diacylglycerol^41^, the conical shape of DIM may modulate membrane protein activity. Accordingly, we find that DIM increase the non-opsonic phagocytosis of zymosan, a process well known to be mediated by a repertoire of membrane receptors, including complement receptor 3 (CR3)^32^ and the mannose receptor^42^. DIM may act on membrane proteins via different biophysical mechanisms. First, DIM may impose curvature on the host membrane^12^, which in turn may modulate integral membrane protein sorting^43, 44^ and function^45^. DIM could also trigger reorganization of lipid nano-domains to modulate signaling platforms^14, 46^. Thus, our findings should open new avenues for understanding how *Mtb* subverts other receptors involved in its recognition by the immune system^47^, including Toll-like receptors, NOD-like receptors, and C-type lectin receptors.

Modulating the activity of membrane proteins will ultimately modulate cellular functions. Indeed, we showed that DIM promote *Mtb* infection (**Fig. 5**). To confirm the relevance of the conical shape, we showed that conical POPE lipids, but not cylindrical POPC lipids, added to macrophages also restored the infection capacity of a DIM-deficient *Mtb* mutant and improved phagocytosis (**Fig. 5**). Our results also shed new light on previous observations showing that a DIM-deficient *Mtb* mutant coated with tripalmitin lipid was less effective in infecting macrophages than DIM-coated mutants^8^. This can now be understood by the fact that tripalmitin did not promote formation of a non-bilayer phase transition in our model membranes (**Fig. 5**), hence did not display a strong conical shape. To our knowledge, our results demonstrate for the first time that the conical shape of a lipid promotes phagocytosis. The conical shape of DIM and their effect on disorganizing the membrane may also play a role in the induction of phagosomal membrane rupture and cell death^48, 49^. Altogether, this new understanding of how the molecular shape of DIM lipids and their biophysical properties affect biological membranes may help to design host-directed therapeutic strategies to fight Tuberculosis^50^ by preventing the infection of macrophages by *Mtb*.

## ACKNOWLEDGEMENTS

This work was supported by the CNRS-MITI grants PEPS MPI 2018 and “Modélisation du vivant” 2019, Agence Nationale de la Recherche (grant ANR-16-CE15-0003), the Fondation pour la Recherche Médicale FRM (“Equipe FRM” number DEQ20160334879) and the Centre National de la Recherche Scientifique (CNRS). J.A. was a recipient of a PhD scholarship from the French government. This work was granted access to the HPC resources of CALMIP supercomputing center under the allocation 2019-17036. C.A.D. thanks TRI-Genotoul Imaging facility (Toulouse, France). We acknowledge Life Science Editors for proofreading the manuscript. We thank Olivier Neyrolles, Jérôme Nigou, Laurence Salomé, and Justin Teissié for fruitful discussions and support.

## METHODS

### Antibodies, lipids and reagents

The rabbit polyclonal antibody against mycobacteria was produced as previously described^8^. The Rhodamine Red-conjugated goat anti-rabbit secondary antibody and Wheat Germ Agglutinin (WGA), Alexa Fluor^®^ 350 conjugate were purchased from Invitrogen. DIM were extracted from *Mycobacterium canetti* as described below. 1-palmitoyl-2-oleoyl-*sn*-glycero-3-phosphocholine (16:0-18:1 PC, POPC), 1-palmitoyl-2-oleoyl-*sn*-glycero-3-phosphoethanolamine (16:0-18:1 PE, POPE), 1-stearoyl-2-oleoyl-*sn*-glycero-3-phosphocholine (18:0-18:1 PC, SOPC), 1,2-dioleoyl-*sn*-glycero-3-phosphoethanolamine (18:1-18:1 PE, DOPE), 1-palmitoyl-2-hydroxy-*sn*-glycero-3-phosphocholine (16:0 lysoPC) and 1,2-dilinoleoyl-sn-glycero-3-phosphoethanolamine (18:2 PE, DLiPE) were purchased from Avanti Polar Lipids (Alabaster, AL, USA). Tripalmitin was a generous gift from M. Tropis (Toulouse, France). The other reagents were purchased from Sigma-Aldrich, except when specifically mentioned.

### Bacterial strains and growth conditions

The strains used in this study included the wild-type (WT) *M. tuberculosis* strain, H37Rv Pasteur (the sequenced strain from Institut Pasteur, Paris) and two distinct H37Rv mutants. The *ppsE* mutant (H37Rv*ΔppsE*) was constructed in a previous study by insertion/deletion within the polyketide synthase gene *ppsE*^51^ required for the synthesis of DIM. The WT strain and the H37Rv*ΔppsE* mutant were rendered fluorescent by the transfer of plasmid pMV361H *gfp*^8^. The *lppX mutant (*H37Rv Δ*lppX)* was constructed by homologous recombination using the thermosensitive counterselectable plasmid pPR27 as previously described ^52^. Briefly, a 2.6kb DNA fragment covering the *lppX* gene was amplified by PCR from H37Rv genomic DNA using primers lppXA (5’-GCTCTAGAGTTTAAACGCATTTGAGCAGCCGAG-3’) and lppXB (5’-GCTCTAGAGTTTAAACGAAGAATACCTGGCCGC-3’) and inserted into a cloning vector. The *res-Ωkm-res* cassette (Malaga et al., 2003) was inserted at the unique KpnI site within the *lppx* gene to generate the allelic exchange substrate (AES) formed of the *res-Ωkm-res* cassette flanked by two arms (of approximately 1kb) specific to *lppX*. This AES was recovered on a PmeI restriction fragment and inserted into the XbaI site of pPR27. The resulting plasmid was transferred into the recipient *M. tuberculosis* H37Rv strain and allelic exchange mutants were selected as described previously^52^. Kanamycin and sucrose resistant clones were analyzed by PCR using primers lppXC (5’-CAAACGCGTTTCTGGACGG-3’), lppXD (5’-GGCAATCCACACGGTCGC-3’), lppXE (5’-GAGCATTGAAAGCTCCCACC-3’) specific of the *M. tuberculosis* H37Rv genome, and res1 (5’-GCTCTAGAGCAACCGTCCGAAATATTATAAA-3’) and res2 (5’-GCTCTAGATCTCATAAAAATGTATCCTAAATCAAATATC-3’) specific of the *res-Ωkm-res* cassette. One clone giving the pattern corresponding to the allelic exchange was retained for analysis and named H37Rv Δ*lppX (or PMM76).* All strains were cultured at 37°C in Middlebrook 7H9 liquid medium (BD Difco) containing 10% albumin-dextrose-catalase (ADC) (BD Difco). When required, kanamycin, hygromycin and Tween-80 were added to the medium to a final concentration of 40 μg mL^−1^, 50 μg mL^−1^ and 0.05% (v/v) respectively.

### Purification of DIM and preparation of lipid solutions

DIM were purified from *M. canetti* as previously described^53^. Briefly, total mycobacterial lipids were extracted from stationary cultures of *M. canetti*. The bacteria were left successively in CH_3_OH/CHCl_3_ (2:1, vol/vol) for 48 h and in CH_3_OH/CHCl_3_ (1:2, vol/vol) for 24 h. The organic phase was recovered, washed with water and dried. Total lipids were then resuspended in CHCl_3_ and the chromatographic separation of DIM was run manually using Sep-Pak Silica Classic Cartridges (55-105 µm particle size; Waters) and an elution gradient of an increasing concentration of diethylether (0-10% (v/v)) in petroleum ether. Fractions containing the isolated compounds were pooled and dried. Stock solutions of purified DIM (40 mg/mL), POPE (20 mg/mL) and POPC (21 mg/mL) were prepared by dissolving the dried lipids in CH_3_OH/CHCl_3_ (2:1, vol/vol). The solutions were then injected in serum-free RPMI 1640 medium (Gibco) at the final concentration of 70 µM (1/400 dilution) and sonicated at 37°C until complete dispersion of the lipids.

### Macrophage culture

The human promonocytic cell line THP-1 (ECACC 88081201; Salisbury, UK) was cultured in RPMI 1640 medium containing 10% heat-inactivated fetal bovine serum (FBS), 2 mM L-Glutamine, 1 mM sodium pyruvate, and 1% MEM non-essential amino acids. For macrophage differentiation, the THP-1 cells were washed and suspended in medium containing 10% FBS. The cells were distributed in a glass petri dish at a density of 3 × 10^6^ cells/petri dish and were differentiated into macrophages with 30 nM phorbol 12-myristate 13-acetate (PMA) for 3 days. Before use, the cells were washed twice with fresh medium.

Human blood purchased from the Etablissement Français du Sang in Toulouse (France) was collected from fully anonymous non-tuberculous donors. Human macrophages derived from monocytes (hMDMs) were prepared as previously described^8^. Briefly, monocytes were isolated from peripheral blood mononuclear cells (PBMC) by adhesion on a glass coverslip in 24-well tissue culture plates. Monocytes (5 × 10^5^ cells/well) were differentiated into hMDMs in RPMI 1640 (Gibco), supplemented with 2 mM glutamine (Gibco) and 7% (v/v) heat-inactivated human AB serum for 7 days.

### Macrophage infection

Single cell suspensions were prepared with exponentially growing strains as previously described^8^. Briefly, the bacteria were grown to mid-exponential growth phase on Middlebrook 7H9 liquid medium supplemented with 10% ADC, and were then pelleted by centrifugation and dispersed in serum-free RPMI 1640 medium using glass beads. The number of bacteria per mL in the suspension was estimated by measurement of the optical density at 600 nm. The bacteria were added to the macrophages at the indicated multiplicity of infection (MOI) and incubated for 1-2 h at 37°C in an atmosphere containing 5% CO_2_. Extracellular bacteria or particles were removed by three successive washes with fresh medium.

### Assay for monitoring DIM transfer to macrophage membranes

For experiments with purified DIM, THP-1 cells were incubated with RPMI 1640 medium supplemented with 70 µM DIM at 37 °C and 5% CO_2_. After 1 h, the cells were rinsed with fresh medium and detached by incubation with a 0.05% trypsin-EDTA solution (Gibco) for 15 min. The cells were then harvested, centrifuged at 150 × g for 10 min and the cell pellet was suspended in RPMI 1640 medium (A).

For experiments with DIM in the context of the *M. tuberculosis* cell envelope, THP-1 cells were incubated with H37Rv WT or H37Rv *D*LppX (MOI 15:1), washed with fresh RPMI-1640 medium, and further incubated in the presence of serum at 37°C and 5% CO_2_. After 40 h, the cells were rinsed with RPMI-1640 medium, detached enzymatically with trypsin and centrifuged at 150 × g for 10 min. The membranes were prepared using a protocol adapted from Rhoades *et al*. ^33^. The pellet was suspended in 1 mL ice-cold homogenization buffer (1 mM EDTA, 20 mM HEPES, pH 7) containing 250 mM sucrose and the cells were disrupted by 25 passages through a 26-gauge needle. Following centrifugation at 3000 × g for 10 min at 4°C in order to sediment nuclei and large cell debris, the supernatant was recovered. This step was repeated twice. The supernatant was layered onto a linear gradient of 30% to 12% sucrose and centrifuged at 2,000 × g for 1 h at 4°C. The upper portion of the gradient containing the membrane fraction was isolated, layered on a discontinuous gradient of 50% to 25% sucrose, and centrifuged at 2,000 × g for 30 min at 4°C. The fraction above the 25% portion containing the membrane fraction was isolated, centrifuged at 110,000 × g for 1 h at 4°C and the membrane pellet was taken up in homogenization buffer (B).

Total lipids were extracted from the macrophage membranes using the Bligh and Dyer extraction protocol^54^. Briefly, to one volume of cells suspension (A) or membrane fraction (B), 2.5 volumes of CH_3_OH and 1.25 volumes of CHCl_3_ were added. The mixture was incubated at room temperature for 48 h. Then, 1.25 volumes of CHCl_3_ followed by 1.25 volumes of CHCl_3_ were added. This mixture was left standing for 24 h to separate the organic and aqueous phases. Twenty-four hours later, the organic phase containing the lipids was recovered and dried under a stream of nitrogen. For the membrane fraction samples, the DIM were purified from the total lipid extract by column chromatography using a Florisil column. The Florisil was equilibrated with a solution of petroleum ether/diethyl ether (98:2, v/v). The total lipid extract was dissolved in this solution and DIM were eluted with the same solution.

### Lipid analysis by MALDI-TOF MS analysis

DIM were analyzed by matrix-assisted laser desorption-ionization time-of-flight (MALDI-TOF) mass spectrometry, as described previously^55^. Lipid residues were dissolved in 20 μL of CHCl_3_, deposited on the analysis plate and dried. Then, 0.5 µL of 2,5-dihydroxybenzoic acid (10 mg/mL) dissolved in CHCl_3_/CH_3_OH (1:1, vol:vol) were deposited on the sample and allowed to crystallize at room temperature. Mass spectra were acquired with a MALDI TOF/TOF 5800 analyzer (Applied Biosystems/AB SCIEX, Framingham, MA, USA) equipped with an Nd:YAG laser (Wavelength 349 nm; pulse rate 400 Hz). The acquisition was carried out in continuous scan mode, in positive mode with a laser intensity of 3500 (arbitrary unit of the software). The final spectrum was obtained by accumulating 10 spectra of 250 laser shots.

### Phagocytosis assay

Phagocytosis was assessed as described previously^8^. Briefly, human monocyte-derived macrophages (hMDM) cultured on sterile glass coverslips in 24-well culture plates were infected with GFP-expressing bacteria (MOI 10:1) or zymosan (MOI 30:1) for 1 h. When indicated, the hMDM were pre-treated for 1 h with 70 µM lipids or 1/400 lipid solvent (CH_3_OH/CHCl_3_ (2:1, vol/vol) prepared in RPMI as for lipid suspension). At the end of infection, the hMDM were intensively washed and fixed with 4% (w/v) PFA. For mycobacteria, extracellular GFP-bacilli were labelled with rabbit anti-mycobacteria Ab revealed by a Rhodamine Red-conjugated goat anti-rabbit secondary Ab. The cells were then permeabilized with 0.3% Triton X-100 for 5 min and stained for 15 min with WGA conjugated to Alexa Fluor^®^ 350 to visualize the hMDM. For zymosan, hMDMs were labelled with TRITC-phalloidin and extracellular zymosan was readily distinguished from internalized particles, which appeared as yellowish grains within a dark phagosome bordered by diffuse red staining. The percentage of cells having ingested at least one bacterium or zymosan particle was determined by fluorescence microscopy using a Leica 43 DM-RB epifluorescence microscope. For each set of conditions, the experiments were performed in duplicate and at least 200 cells were counted per slide. Data are presented as the mean ± standard error of the mean (SEM) of the indicated number of experiments (n). Data were analyzed by the Wilcoxon signed-rank test using GraphPad PRISM (GraphPad Software, GPW5-078069-NBH9780) and p < 0.05 was used as the limit of statistical significance.

### Formation of multilamellar vesicles for HRMAS NMR experiments

To form multilamellar vesicles (MLV) of well-defined lipid compositions, we mixed appropriate volumes of chloroformic stock solutions of the different lipids in glass tubes. The chloroform was evaporated using a rotating evaporator. We inclined the tubes in the evaporator, resulting in the formation of a thin lipid film over a large area of the tube. The lipids were further dried under vacuum for ∼2 h to remove any remaining traces of the solvent. Next, we added sufficient Tris buffer (10 mM Tris, 1 mM EDTA, pH 7.4) to the tubes to cover the lipid film. The lipids were left to hydrate at a temperature chosen to favor the formation of the lamellar phase of the liposome membranes (see below for further details). We then vortexed each tube 6 × 30 s. After vortexing, we obtained a cloudy suspension of liposomes. We transferred the liposome suspensions to 1.5 mL centrifuge tubes and centrifuged the tubes for 15 min at 16000 g, then stored them at 4°C. Before each NMR experiment, we removed the supernatant and transferred 50 µL of the liposome pellet to a 4 mm MAS rotor, taking care not to warm the liposomes. Before introducing the rotor into the NMR spectrometer, we equilibrated the temperature to ∼5°C.

In order to be sure to prepare the membranes in the lamellar phase, we adapted the temperature of hydration to the lipid composition. Pure DOPE has a lamellar-to-inverted-hexagonal transition temperature of 10°C. Accordingly, we hydrated the samples of DOPE, DOPE/SOPC (9:1) and DOPE/SOPC (5:1) for 15 min on ice. We vortexed the lipids 6 × 30 s (with cooling on ice for at least 30 s between vortexing periods) in order to prepare and conserve the liposomes in the lamellar phase. DOPE/SOPC (3:1), DOPE/SOPC (1:1) and pure SOPC (which do not form a H_II_ phase at 30°C) were incubated at 30°C in a water bath and vortexed 6 × 30 s with return to the water bath.

Incorporating the highly hydrophobic lipid DIM into liposomes requires incubating the lipids at higher temperature, but this would have favored the formation of the inverted-hexagonal phase for some membrane compositions. We therefore adapted our protocol for DIM-containing liposomes and their controls without DIM. For these lipid mixtures, we hydrated the lipids at 37°C on a shaker overnight. We then transferred the tubes to a water bath set at 37°C. The tubes were vortexed for 6 × 30 s, with return to the water bath between vortexing periods. All liposomes with compositions DOPE/SOPC (3:1) +/− DIM, DOPE/SOPC (3:1) +/− TAG and DOPE/SOPC/lysoPC (75:20:5) +/− DIM were prepared according to this protocol. We checked the correct incorporation of DIM in the liposomes using thin-layer chromatography on a few µL of the liposome suspension. We also conducted a separate ^1^H-NMR experiment to quantify the incorporation of DIM in liposomes using our protocol (see Supplementary Material and Figures).

### NMR data acquisition

Phosphorus NMR spectra were acquired on a 500 MHz Bruker Avance spectrometer, in a HRMAS probe, with deuterium lock. The lipid samples (typically 6 mg total lipids in 50 µL of Tris buffer (10 mM Tris, 1 mM EDTA, pH 7.4) were inserted into 4 mm rotors with spherical inserts. The temperature of the sample could be varied between 278 K and 324 K and was controlled to ± 0.1 K with a Bruker variable temperature unit. The temperature was calibrated using the known temperature dependence of methanol chemical shifts. 31P chemical shift anisotropies (CSA) were determined from the spinning sideband manifolds at a spinning frequency of 2000 ± 1 Hz. MAS spectra were obtained with a spin-echo sequence (π/2 - τ - π) where the π/2 pulse had a length of 5.3 µs (at a power of 107 W), applied at the lipids’ isotropic resonance frequency, and the interpulse delay, τ, was 20 µs. The dwell time was 1 µs, the acquisition time 65 ms, the relaxation delay 1 s and the number of scans 4096. No proton decoupling was applied during acquisition, since it was shown to have no effect on the linewidth at a 2 kHz spinning frequency (i.e. well above the 1H-31P dipolar coupling of ∼500 Hz in fluid lipid bilayers). For every sample, a ^1^H NMR spectrum at a spinning frequency of 10 kHz was acquired in order to calibrate the ^1^H (using the methylene peak at 1.25 ppm with respect to TMS) and 31P chemical shifts (with respect to phosphoric acid at 0 ppm, using *γ*_P_/*γ*_H_ = 0.40480742). The observed ^31^P isotropic chemicals shifts were −1.00 ± 0.02 ppm for the phosphatidylcholine head group (in SOPC) and *−*0.27 ± 0.02 ppm for the phosphatidylethanolamine head group (in DOPE) and varied only slightly with the lipid compositions explored. Typical linewidths were 40-60 Hz (i.e. ∼0.25ppm) so that the PC and PE sideband patterns were well resolved and could be fitted independently.

Every liposome sample was equilibrated overnight at 277 K before NMR measurements. We then measured its ^31^P MAS spectrum at temperatures from 284 K to 324 K, in 2 K increments every 100 min (25 min equilibration plus 75 min acquisition time for each temperature). For lipid mixtures which did not undergo a lamellar to inverse-hexagonal phase transition (i.e. with a small proportion of DOPE) the spectra were shown to be fully reversible when going down from 324 K to 284 K. However, once the H_II_ phase had formed, it did not revert to the lamellar phase, at least after one day of equilibration at 284 K. Hence, all measurements were done for increasing temperatures, starting from the lowest temperature, taking care of maintaining the liposomes at low temperature during their preparation and before NMR measurements. The actual insertion yield of DIM in the lipid bilayers was determined by ^1^H-NMR (see Supplementary material).

### Spectral deconvolution

We analyzed each ^31^P spinning sideband manifold using the solid line shape analysis tool (SOLA) available in Topspin 3.5. CSA parameters were calculated using the Haeberlin convention for the anisotropy values Δδ = δ// − δ⊥ as commonly done in the membrane literature and as explained before^56^. We estimated the uncertainty of Δδ values to be ± 0.2 ppm, based on several measurements and fitting performed on independent samples. We first determined the CSA parameters of lipids organized in a single phase (e.g. SOPC for the lamellar Lα phase and DOPE for the HII phase), at several temperatures between 284 K and 324 K. The CSA parameters for each lipid and each phase varied slightly and linearly with temperature, due to increased motion at higher temperatures: from 49.6 ± 0.2 ppm (293 K) to 45.0 ± 0.2 ppm (333 K) for SOPC in the Lα phase; from *−*22.2 ± 0.2 ppm (278 K) to *−*20.1 ± 0.2 ppm (333 K) for DOPE in the HII phase. The following linear regressions were obtained: Δδ = *−*0.0944 T + 76.599 for SOPC in the Lα phase, Δδ = *−*0.0409 T *−*33.627 for DOPE in the HII phase, where T is the temperature in Kelvin; these regressions were used to calculate CSA values at intermediate temperatures. Since the lipids in the HII phase experience additional motional averaging due to lateral diffusion around the aqueous channels, their CSA is obtained by averaging two components δ_//_ and δ_⊥_, divided by two, for reasons discussed by Cullis and De Kruijff^57^. The CSA values in the L*_α_* and H_II_ phases have opposite signs. Hence, the CSA value of SOPC in the H_II_ phase was taken to be *−*0.5 times its value in the L_α_ phase, while the CSA value of DOPE in the L*_α_* phase was 2 times its value in the H_II_ phase. Knowing the temperature-dependent CSA parameters for both lipids in both phases allowed a drastic reduction of the number of parameters to be fitted when analyzing lipid mixtures in which the two phases coexist. After optimizing the peak positions and linewidths, the SOLA fitting algorithm only has to search for the L*_α_*-H_II_ proportions that best reproduce the experimental spectra. The intensity of the spinning sideband at +4 kHz is a good reporter of the proportion of Lα phase, since this peak is almost absent from the typical H_II_ phase spinning sideband manifold^58^ (see Fig. 4c). This fitting protocol was very robust and prevented the artefacts and instabilities arising from fitting too many parameters. We should stress that our procedure is based on the assumption that the spectra consist of a linear combination of the spectra of the two lipids in the L*_α_* and H_II_ phases, possessing CSA parameters identical to their values in the pure phases at the same temperature. This simplifying assumption was sufficient to characterize the lipid phase transitions, although we cannot exclude that the actual lipid behavior is more complex.

### Quantum mechanics

To calculate partial charges for DIM lipid, a molecular dynamics simulation was first performed for 40 ns in the NPT ensemble. Lipid14^59^ and general AMBER force field (Gaff)^60^ were used in order to describe bonded and non-bonded terms within the AMBER program. From this trajectory, 30 structures were extracted based on the clustering method (*kclust* from the MMTSB Toolkit, http://www.mmtsb.org, with the radius of 2.5 Å). These structures were fully optimized at the HF/6-31G* quantum chemical method using the Gaussian 09 suite of programs (http://gaussian.com/). Partial charges were then obtained using RESP method^61^ (RESP-A1) implemented in the RED tools^62^. For each structure, partial charges where obtained by charge fitting to the electrostatic potential at points selected according to the Merz-Singh-Kollman scheme^63, 64^ as proposed by the RESP-A1 method. Before charge calculations, a reorientation procedure was applied in order to maintain one ester (O=C-O) group in the same orientation for all 30 structures. Partial charges for the head groups of the DIM model are displayed in Figure S2.

### MD simulations

The atomistic MD simulations were performed with the Amber16 software^65^ (http://ambermd.org). The system contained a membrane of 300 POPC molecules and was solvated with TIP3P water model using the CHARMM-GUI server^66, 67^. The DIM molecule was positioned in the POPC bilayer with the polar core in the proximity of the POPC oxygen atoms (see Fig. 2a). In order to avoid steric clashes of the DIM molecule with POPC, the system was initially minimized by executing 1500 iterations of the steepest descent (SD) algorithm, followed by 1500 iterations of the conjugate gradient (CG) algorithm, with weakly restrained solute (k = 10 kcal/mol/Å^2^). Next, a short 100 ps MD run was performed on weakly restrained solute with temperature varying linearly from 0 to 303 K. The temperature control was achieved using the Langevin dynamics with the collision frequency parameter γ equal to 1.0 ps^−1^. The integration step used in this run was 1 fs. Throughout the calculations a cutoff of 10 Å was used for electrostatic interactions. The MD simulation continued with the equilibration of the system, consisting of 10 consecutive MD steps of 500 ps each, at constant temperature of 303 K with no restraints, with the integration step of 2 fs. The production run was then launched for 800 ns with constant pressure of 1 bar. The Langevin dynamics was used to control the temperature, with γ = 1.0 ps^−1^, while the pressure was controlled by the anisotropic Berendsen barostat with the pressure relaxation time τ_p_ = 1 ps. Bonds involving hydrogen were constrained with the SHAKE algorithm.

Coarse-grain simulations were performed using GROMACS 2016^68^ with the MARTINI force field 2.2^69, 70^. We have designed the CG model of DIM by following the original MARTINI parametrization strategy^71^. We approximated 3-4 heavy atoms constituting the long hydrophobic acyl chains by one C1 particle. Small methyl branched parts were approximated by a small particle SC1. The slightly polar extremity of the phthiocerol chain was approximated by a N_0_ particle while the glycerol ester moieties were represented by an intermediate hydrophobicity particle Na as usually done for other phospholipids (see Fig. 2b). Force constants and equilibrium values for bonds and angles were extracted from the atomistic simulation (see Fig. S3). All the systems were first gradually equilibrated as proposed in the CHARMM-GUI MARTINI Maker protocol^72^). Coulomb interactions were treated using the reaction-field potential and Lennard–Jones interactions were treated using shifted potentials with a cut-off radius of 1.1 nm. For systems with a pure SOPC bilayer or DIM molecules embedded in POPC bilayer, pressure was maintained at 1 bar using the Parrinello-Rahman algorithm^73^ with a semi-isotropic pressure control. The temperature was kept at 310 K using the v-rescale algorithm^74^. For pure DOPE and SOPC-DOPE systems undergoing lamellar to hexagonal transitions, we used the protocol described by S.J. Marrink and A. E. Mark^27^. We stacked 4 membranes of SOPC-DOPE mixture at a 3:1 ratio with a hydration level of approximately 2-3 water CG-particles per lipid (equivalent to c.a. 9-10 atomistic water molecules) in order to see the phase transition in a reasonable amount of time as detailed in S.J. Marrink and A. E. Mark paper^27^. A Berendsen thermostat in combination with a Berendsen barostat^75^ were used. A fully anisotropic coupling pressure was applied with a reference pressure of 1 bar. Systems were equilibrated at 280K and then 8 different simulations were launched with temperatures ranging from 280K to 350K with an increment of 10K. In all CG systems, a time step of 20 fs was used. See Table S1 for a summary of the simulations. Parameters for CG and atomistic representations of DIM lipids will be available on the MARTINI website: cgmartini.nl. Density profiles were performed using the Gromacs tool density (http://manual.gromacs.org/documentation/2016/onlinehelp/gmx-density.html). MD simulations figures were performed using VMD^76^. Scripts used to analyze MD simulations will be available at: https://github.com/MChavent

