## Supplementary Material for "The conical shape of DIM lipids promotes *Mycobacterium tuberculosis* infection of macrophages"

### Supplementary Material and Figures

#### Tilt angle analysis and curve fitting

The lamellar to inverted hexagonal phase transition was characterized from the coarse-grained (CG) simulations data by analyzing the distributions of the local orientations of SOPC and DOPE. The orientations of these molecules are described by the tilt angle $\theta$ between their director (*i.e*. the principal axis of the inertia tensor) and the $z$-axis of the CG simulation space (aligned with the membrane normal in the lamellar phase). When $\theta > \pi/2$, we use $\theta' = \pi - \theta$ to uniformly describe the lipid orientation for the two membrane leaflets. Lipid orientation distributions were obtained by counting molecules in 100 bins of width $\delta\theta = \pi/200$ from $0$ to $\pi/2$ for the last 250 frames of the CG simulations of pure SOPC at 310 K, pure DOPE at 350 K, and DOPE/SOPC/DIM mixtures at several temperatures, except with 2.5% DIM at 340 K where the last 125 frames were used (due to a late transition). The distributions were then normalized by their integral as evaluated from molecule and frame numbers.

<u>Lamellar phase:</u> In the lamellar phase, due to periodic boundary conditions, the bilayers are globally parallel to the $(xOy)$ plane and normal to the $(Oz)$ axis. The director of each lipid is on average parallel to $(Oz)$. The local orientation is expressed by the lipid's polar angle $\theta \in [0, \pi/2]$. We assume that the deviation from the normal orientation is described by a simple elastic model with an energy given by $E(\theta) = -\kappa_1 \cos\theta$, where $\kappa_1$ is a bending elastic modulus, similar to an electric dipole in an external field. With this expression for the energy $E(\theta)$, the equilibrium Boltzmann distribution of the angle $\theta$ at temperature $T$ is given by the equation:

$$p_{\mathrm{lam}}(\theta) = \frac{1}{Z} \sin\theta \, e^{\tilde{\kappa}_1 \cos\theta} \quad \text{(Eq. S1)}$$

where $Z = (e^{\tilde{\kappa}_1} - 1)/\tilde{\kappa}_1$ is the normalization factor and $\tilde{\kappa}_1 = \kappa_1/(k_B T)$, $k_B T$ being the thermal energy. The reference distribution function of the lipid orientation in the lamellar phase was obtained by fitting CG data of SOPC at 310 K with Eq. S1, as illustrated in the figure below where the fitted value is $\tilde{\kappa}_1 \simeq 12$.

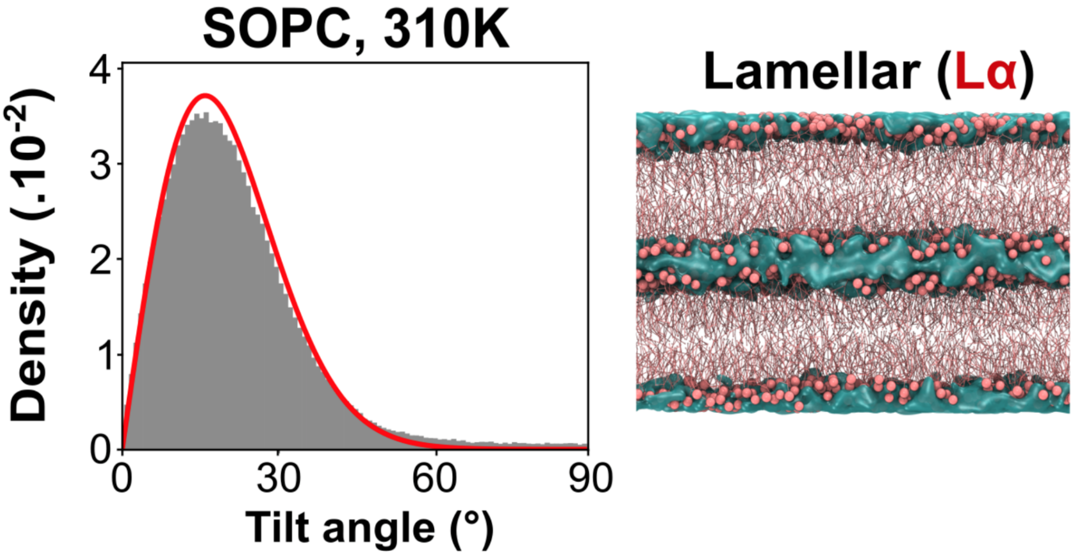


**Distribution of lipid orientations in the lamellar phase.** Left: Histogram (in gray) of tilt angles θ extracted from CG simulations. The red curve is a fit to the probability distribution $p_{lam}(\theta)$ (Eq. S1) given in the text with $\tilde{\kappa}_1 \simeq 12$. Right: Lamellar SOPC system used for the fit.

When the lamellar and hexagonal phases coexist, as discussed in the main text, the bilayers are not as regular as in the pure lamellar phase. One type of possible perturbation is that the bilayers are tilted with respect to the normal orientation $\theta = 0$, by an average angle $\theta_0 > 0$. To simulate this effect, angles $\theta$ were sampled as follows (see figure below, upper left): a fixed vector making an angle $\theta_0$ with $(Oz)$ is defined and sampled lipid directors make an angle $\theta'$ with it following the same distribution as in equation S1. The angles $\theta$ that these lipid directors make with $(Oz)$ are then calculated. We have tested a simple phenomenological fitting law:

$$p_{\text{lam},2}(\theta) = \frac{\beta}{Z} \sin\theta \, e^{\kappa_1 \cos(\theta - \theta_0)/k_B T} \qquad \text{(Eq. S2)}$$

that gives very satisfactory fits (data not shown), but with a fitted parameter $\theta_{0,\text{fit}}$ different from the original value $\theta_0$, as shown in the table below. Note that, in addition, the normalization factor $1/Z$ must be rescaled by a prefactor $\beta$ to ensure that the total probability remains equal to 1.

| $\theta_0$ (°) | 10 | 15 | 20 | 24 | 30 | 40 |
|---|---|---|---|---|---|---|
| $\theta_{0,\text{fit}}$ (°) | 4 | 8 | 13 | 19 | 26 | 37 |
| $\beta$ | 0.75 | 0.57 | 0.42 | 0.33 | 0.26 | 0.19 |

**Fitting parameters table.** Values of the fitted parameters $\theta_{0,\text{fit}}$ and $\beta$ for given tilt angles $\theta_0$ for $\tilde{\kappa}_1 \simeq 12$. Angles are given in degrees.

When fitting the simulated data, one measures the angle $\theta_{0,\text{fit}}$. From its value, the prefactor $\beta$ must be inferred. For this, we propose the following interpolation function, based on the values in the table above for $\tilde{\kappa}_1 = 12$:

$$\beta(\theta_{0,\text{fit}}) \simeq 0.67e^{-0.11\theta_{0,\text{fit}}} + 0.33e^{-0.015\theta_{0,\text{fit}}} \qquad \text{(Eq. S3)}$$

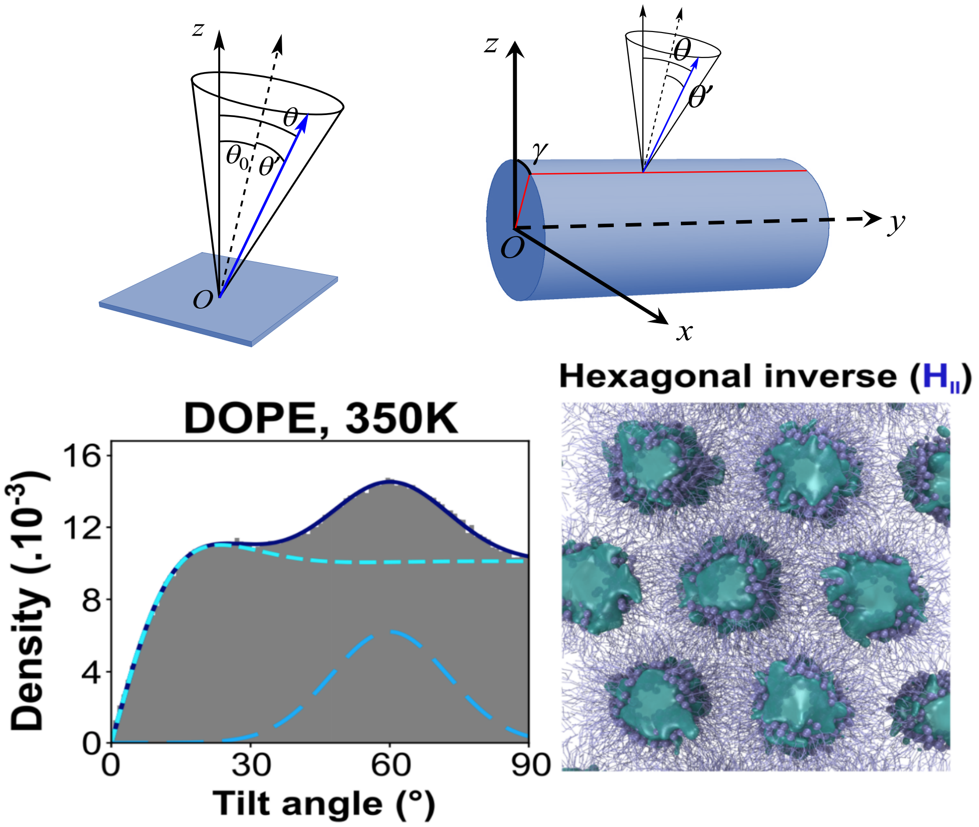


**Distribution of lipid orientations in the inverted hexagonal phase.** Upper left: In the lamellar phase case, construction of a lipid director (blue arrow) isotropically distributed around a fixed tilt direction with polar angle $\theta_0$ (dashed arrow). The membrane is represented by the blue plane parallel to $(xOy)$. Upper right: The hexagonal phase is made of parallel lipid cylinders (in blue) with axes chosen here to be parallel to the $(Oy)$ axis, without loss of generality. For a given lipid, its director is on average normal to the cylinder surface (dotted arrow). It is characterized by the polar angle $\gamma \in [0,2\pi[$. The lipid director (blue arrow) is isotropically distributed around this average direction. Lower left: Histogram (in gray) of tilt angles $\theta$ (in degrees) from MARTINI simulations. The dark blue solid curve is a fit with the probability distribution $p_{\text{hex}}(\theta)$ (Eq. S2) given in the text. The cyan dashed line is the probability distribution $p_{\text{hex},0}(\theta)$ if the hexagonal phase were ideal. The blue long-dashed line represents favored areas for lipid tails in between the water tubes (see text below). Lower right: hexagonal DOPE system used for the fitting.

Inverted hexagonal phase: An ideal inverted hexagonal phase is made of parallel cylinders whose axes form a hexagonal array. If the distance between these cylinders is sufficiently large, the orientation of lipids is isotropic around each cylinder. As illustrated in the figure above (upper right), the average orientation of a lipid director is perpendicular to the cylinder surface. We denote by $\gamma$ the corresponding polar angle, as displayed in the figure. As in the lamellar phase, we assume that due to thermal fluctuations, lipid directors actually make an angle $\theta'$ with this average orientation with a probability distribution $p(\theta') = \frac{1}{Z} \sin\theta' \, e^{\tilde{\kappa}\cos\theta'}$. We again sampled angles $\theta$ by drawing a random angle $\gamma$ uniformly on $[0,2\pi[$ and a random angle $\theta'$. We successfully fitted (not shown) the thus obtained data with the following heuristic law, where $Z_{\text{hex},0}$ is a normalization factor:

$$p_{\text{hex},0}(\theta) = \frac{1}{Z_{\text{hex},0}}\left[1 - e^{-\frac{\theta}{\tilde{\kappa}_{21}}} \cos\left(\frac{\theta}{\tilde{\kappa}_{22}}\right)\right] \quad \text{(Eq. S4)}$$

When the distance between cylinders decreases, the lipid tails in opposing cylinders of the inverted hexagonal phase $H_{II}$ repel each other. Some director orientations are favored in coherence with the hexagonal symmetry perpendicularly to the cylinder axes (as already seen by others[1]). To mimic this fact, we again propose a heuristic law by adding a Gaussian distribution to the previous one:

$$p_{\text{hex}}(\theta) = \frac{(1-c)}{Z_{\text{hex},0}}\left[1 - e^{-\frac{\theta}{\tilde{\kappa}_{21}}} \cos\left(\frac{\theta}{\tilde{\kappa}_{22}}\right)\right] + c\,\frac{1}{\sqrt{2\pi}\sigma} \exp - \left[\frac{(\theta-\theta_1)^2}{2\sigma^2}\right] \quad \text{(Eq. S5)}$$

where $0 \leq c \leq 1$ is the weight of the Gaussian correction, which has a mean value $\theta_1 = \pi/3$ and a standard deviation $\sigma$. The reference orientation distribution function of the inverted hexagonal phase was obtained by fitting CG data of DOPE at 350K with Eq. S5. Due to the important number of parameters to be fitted in this equation, the fits were performed with an program developed in-house for stochastic global optimization by simulated annealing (GOSA)[2]. This method allows improving the quality of the fit in the case of a high dimensionality of the parameters space, compared to deterministic gradient-based algorithms such as steepest descent or conjugate gradients. Fitting of the tilt angles in the inverted hexagonal phase (lower left figure) illustrates how well this heuristic law fits the distribution extracted from CG simulations.

Coexistence of lamellar and inverted hexagonal phase:

The distribution of lipid orientations for the DOPE/SOPC/DIM mixture in the CG simulations were fitted with a linear combination of the reference distribution functions given previously (Eq. S1 and S5):

$$p_{mix}(\theta) = A\left[\frac{\beta(\theta_0)}{Z}\sin\theta\, e^{\tilde{\kappa}_1\cos(\theta-\theta_0)}\right] + (1-A)\left[\frac{(1-c)}{Z_{\text{hex},0}}\left[1 - e^{-\frac{\theta}{\tilde{\kappa}_{21}}}\cos\left(\frac{\theta}{\tilde{\kappa}_{22}}\right)\right] + c\,\frac{1}{\sqrt{2\pi}\sigma}\exp - \left[\frac{(\theta-\theta_1)^2}{2\sigma^2}\right]\right] \quad \text{(Eq. S6)}$$

by using GraphPad Prism 5.04 (GraphPad Software, La Jolla California USA, www.graphpad.com). As described above, the tilt angle $\theta_0$ and the rescaled prefactor $\beta(\theta_0)$ were introduced because of perturbations due to phase coexistence. It permits to determine the percentage, $A$, of the lamellar phase contribution for DOPE and SOPC lipids separately in each CG simulation using the values of $\tilde{\kappa}_1, c, Z_{\text{hex},0}, \tilde{\kappa}_{21}, \tilde{\kappa}_{22}$ and $\sigma$ resulting of fitting the reference functions for the lamellar and inverted hexagonal phase. See Figure below for an illustration of the fitting for a tilt angle distribution of DOPE.

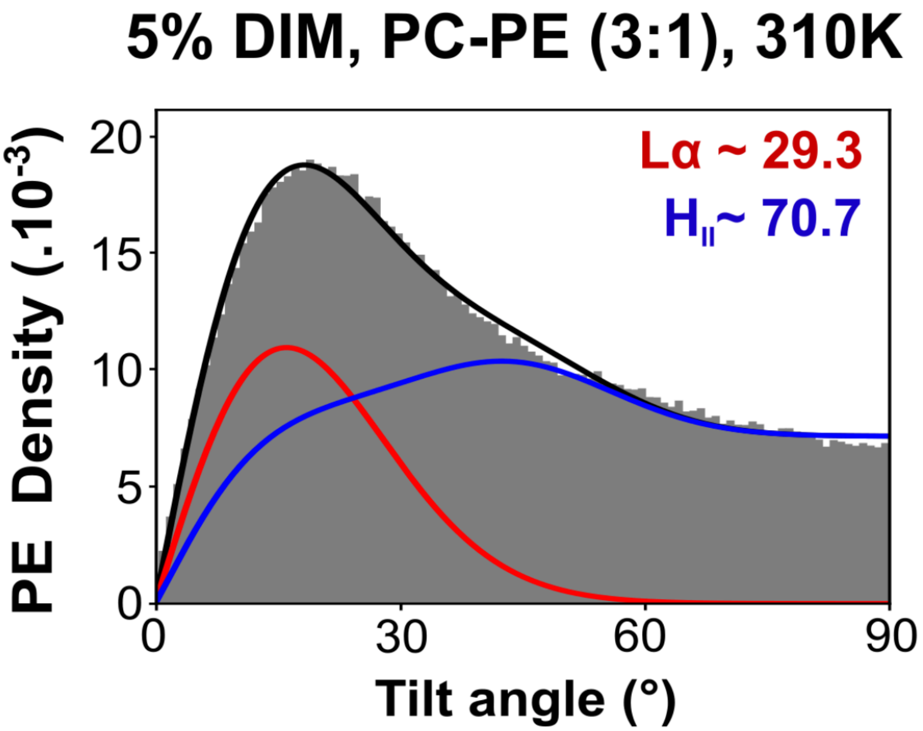


**Distribution of lipid orientations in a mixed-phase system.** DOPE tilt angle distribution for a system containing both inverted hexagonal and lamellar phases (see Fig. 3b). Fitting of the complete distribution (Eq. S6) in black. Individual contributions of lamellar and inverted hexagonal phases in red and blue, respectively.

Phase transition curves were finally obtained by fitting a sigmoid function to the variation of $A$ as a function of temperature for each DOPE/SOPC/DIM mixture using Eq. S7 where $A_0$ and $T_{50}$ respectively stand for the initial lamellar phase contribution and the phase transition midpoint temperature, while $k$ is linked to the slope of the function at $T_{50}$. The same equation was used to fit phase transitions monitored by $^{31}$P NMR.

$$A(T) = A_0 \frac{1}{1+\exp[-k\,(T - T_{50})]} \qquad \text{(Eq. S7)}$$

**DIM quantification in liposomes**

To assess the quantity of DIM incorporated in freshly prepared liposomes, and whether this concentration stayed stable over time, we measured the percentage of DIM actually inserted into liposomes prepared with different percentages of DIM, and equilibrated for several durations and temperature.

Using the protocol for the formation of multilamellar vesicles (see Methods), we formed two batches of liposomes of DOPE and SOPC (3:1) containing 1%, 2.5% and 5% (mol/mol) of DIM, respectively. Each liposome batch contained 100 µg of DIM and appropriate masses of DOPE and SOPC for the targeted DIM percentage. We split each liposome batch into two equal volumes, which we transferred into plastic microcentrifuge tubes (Eppendorf). For each batch, we immediately recovered the liposomes from one tube; we stored the other tube for 24 h at 37°C before recovering the liposomes. To recover the liposomes, we transferred the contents of each tube into a clean microcentrifuge tube. We centrifuged the tube for 15 min at 16000 g, discarded the supernatant and resuspended the pellet in 500 µL of ultrapure water (Milli-Q, Millipore). We then transferred the liposome suspension to a clean glass tube. The lipid suspensions were frozen in liquid $N_2$ and freeze-dried for 24 h.

We dissolved the dried lipids in 250 µL of $CDCl_3$ and transferred the lipid solutions into ∅3 mm NMR tubes. We measured the liquid-state $^1$H NMR spectra of all six tubes on a Bruker 600 MHz spectrometer equipped with a TCI cryoprobe. To measure the DIM/phospholipid molar ratio for each condition, we identified characteristic well-separated peaks in the NMR spectra corresponding to DIM and to the phospholipids. For DIM, we used the quintuplet at 4.84 ppm corresponding to a DIM methine proton. For the phospholipids, we used peaks around 5.34 ppm corresponding to the ethylene protons in the DOPE and SOPC unsaturated side chains. We integrated these peaks and compared the areas to calculate the percentage of DIM in the lipids mixtures. The results of this quantification are shown in the table below. We conclude that, within experimental errors, the measured percentages of DIM agree with the calculated percentages, indicating a good incorporation of DIM into the liposomes prepared following our protocol. Moreover, we did not observe a decrease in the percentage of DIM after 24 h incubation, indicating that there is no exclusion of DIM from the liposome membranes.

**Quantification by $^{1}$H-NMR of the percentage of DIM in freshly prepared liposomes and in liposomes stored for 24 h at 37°C.**

| Target percentage of DIM | Measured percentage of DIM | |
|---|---|---|
| | 0 h | 24 h, 37°C |
| 1 % | 0.81 % | 0.89 % |
| 2.5 % | 2.8 % | 2.2 % |
| 5 % | 5.2 % | 5.5 % |

#### Determination of lipid phase transition temperatures by $^{31}$P NMR

Phosphorus NMR has been recognized for a long time as an excellent technique for identifying non-lamellar phases in lipid bilayers[3] and characterizing the dynamics of the phospholipid head groups [4]. In every lipid phase, the phosphate nucleus is subject to specific movements that average its static chemical shift anisotropy (CSA) tensor in a specific way, thus producing a characteristic spectrum. The $^{31}$P CSA of a liposome preparation can be measured in a static probe from its powder spectrum. Under magic angle spinning at a rate slower than the CSA width, the powder spectrum splits into spinning sidebands, from which the CSA parameters can be extracted[5]. This approach has several advantages: it increases the resolution and sensitivity of the experiment, and allows analysing several CSA's in a lipid mixture, as long as the phosphorus atoms in the different head groups have distinct isotropic chemical shifts, which is the case for SOPC and DOPE.

Several lipid mixtures containing varying DOPE:SOPC molar ratios (1:0, 9:1, 5:1, 3:1, 1:1, 0:1) were investigated (see Table S2). As expected, decreasing the proportion of DOPE increased the lamellar ($L_\alpha$) to inverted-hexagonal ($H_{II}$) phase transition temperature[6]. Pure DOPE is already in the $H_{II}$ phase at 5°C, the 9:1 and 5:1 mixtures have midpoint transitions at 24°C and 39°C, respectively, while the 3:1, 1:1 and 0:1 DOPE:SOPC mixtures do not transit to the $H_{II}$ phase below 51°C, the highest temperature tested. In order to investigate the tendency of an extraneous lipid to induce a $L_\alpha$ to $H_{II}$ transition, we focussed on a DOPE to SOPC ratio of 3:1 since the lipid membrane remains in the lamellar phase for this composition within the 9°C to 51°C temperature range, but is on the verge of a transition to the $H_{II}$ phase. With this selected 3:1 ratio, we tested the effect of adding small amounts of lipids on the $L_\alpha$ to $H_{II}$ phase transition: DIM (at 1 mol%, 2.5 mol% and 5 mol%), tripalmitin (at 2.5 mol% and 5 mol%), dilinoleyl-phosphatidylethanolamine (DLiPE, at 5 mol%). We measured $^{31}P$ NMR spectra every two degrees between 282 K and 324 K and deconvoluted these spectra (see Methods section) to obtain the proportion of lipids in the $L_\alpha$ and $H_{II}$ phase. This analysis was performed both on the PC and the PE peak so that the percentage of the two phases could be obtained independently for these two lipids. At each temperature, the PE peak displayed a higher amount of $H_{II}$ phase than the PC peak in the same lipid preparation, in agreement with the propensity of DOPE molecules to favor and to accumulate in the $H_{II}$ phase. Table S2 shows a subset of the spectra that have been acquired and analysed.

**Table S1 | Summary of Molecular Dynamics simulations**

| Systems | Granularity | Number of Particles | Duration (µs) |
|---|---|---|---|
| 1 DIM | AT | 294 | 0.04 |
| 1 DIM + 300 POPC + Waters | AT | 79977 | 0.8 |
| 1 DIM + 400 POPC + Waters<br>(DIM parametrization) | CG | 13450 | 3 |
| 4 DIM + 400 POPC + Waters (~1% DIM) | CG | 13511 | 3 |
| 8 DIM + 400 POPC + Waters (~2% DIM) | CG | 13502 | 3 |
| 16 DIM + 400 POPC + Waters (3.8% DIM) | CG | 13521 | 3 |
| 32 DIM + 400 POPC + Waters (7.4% DIM) | CG | 13491 | 3 |
| 1248 SOPC + Waters (SOPC 310K, Lamellar) | CG | 17982 | 3 |
| 1248 DOPE + Waters (DOPE 350K, Hexa.) | CG | 16682 | 3 |
| 616 SOPC+ 1880 DOPE + Waters<br>(SOPC-DOPE 3:1 280K to 350K) | CG | 37584 | 3x8 |
| 64 DIM + 616 SOPC+ 1880 DOPE + Waters<br>(2.5% DIM SOPC-DOPE 3:1 280K to 350K) | CG | 37320 | 3x8 |
| 128 DIM + 616 SOPC+ 1880 DOPE + Waters<br>(5% DIM SOPC-DOPE 3:1 280K to 350K) | CG | 38856 | 3x8 |
| 64 DIM + 312 SOPC+ 304 LysoPC + 1880 DOPE + Waters<br>(2.5% DIM SOPC-DOPE 3:1 + 10% LysoPC 280K to 350K) | CG | 36104 | 3x8 |

217

**Table S2 | $^{31}$P NMR parameters obtained on several lipids and lipid mixtures at various temperatures.** The spectra were acquired on fully hydrated multilamellar vesicles, at 2 kHz spinning rate. The CSA parameter $\Delta\delta = \delta_{//} - \delta_{\perp}$ was calculated from the spinning sideband manifold, using the solid line shape analysis tool of Topspin 3.5. The $^{31}$P isotropic chemicals shifts were – 1.00 ± 0.02 ppm for the phosphatidyl choline head group (in SOPC) and – 0.27 ± 0.02 ppm for the phosphatidyl ethanolamine head group (in DOPE).

| Sample | Temperature, K | Phase | % Phase | $\Delta\delta$ (±0.2ppm) | Linewidth (Hz) |
|---|---|---|---|---|---|
| POPC | 278 | Lα | 100 | 50,6 | 57 |
| | 293 | Lα | 100 | 48,2 | 50 |
| | 313 | Lα | 100 | 47,3 | 45 |
| SOPC | 293 | Lα | 100 | 49,6 | 50 |
| | 303 | Lα | 100 | 48,8 | 45 |
| | 313 | Lα | 100 | 46,7 | 59 |
| | 333 | Lα | 100 | 45,0 | 60 |
| DOPE | 278 | HII | 100 | -22,2 | 131 |
| | 288 | HII | 100 | -22,1 | 90 |
| | 293 | HII | 100 | -22,1 | 85 |
| | 303 | HII | 100 | -21,3 | 60 |
| | 313 | HII | 100 | -21,2 | 55 |
| | 323 | HII | 100 | -20,1 | 53 |
| | 333 | HII | 100 | -20,1 | 50 |
| DOPE SOPC 9:1 | 293 | Lα PC | 73 | 48,9 | 45 |
| | | Lα PE | 59 | 43,3 | 45 |
| | | HII PC | 27 | -24,5 | 45 |
| | | HII PE | 41 | -21,6 | 45 |
| DOPE SOPC 5:1 | 293 | Lα PC | 100 | 48,9 | 45 |
| | | Lα PE | 87 | 43,3 | 45 |
| | | HII PC | 0 | -24,5 | 45 |
| | | HII PE | 13 | -21,6 | 45 |
| DOPE SOPC 3:1 | 293 | Lα PC | 90 | 48,9 | 45 |
| | | Lα PE | 82 | 43,3 | 45 |
| | | HII PC | 10 | -24,5 | 45 |
| | | HII PE | 18 | -21,6 | 45 |
| DOPE SOPC 1:1 | 293 | Lα PC | 96 | 48,9 | 40 |
| | | Lα PE | 78 | 43,3 | 40 |
| | | HII PC | 4 | -24,5 | 40 |
| | | HII PE | 22 | -21,6 | 40 |

**Table S3 | Lamellar to inverted hexagonal phase transition fitting from $^{31}$P NMR data and CG simulations.** As described in the Supplementary Material section, both NMR and CG-derived temperature-dependent data have been fitted by a sigmoidal phase transition function in order to obtain $T_{50}$ as well as $A_0$ and k. $T_{50}$ is the phase transition midpoint temperature, $A_0$ the initial lamellar phase contribution and k is proportional to the slope of the function at $T_{50}$. For each tested lipid composition, their values are indicated together with their 95% confidence intervals ($CI_{95\%}$) as determined from the fitting procedure. Coefficient of determination ($R^2$), degrees of freedom and standard deviation of the residuals (Sy.x) are also indicated for each fitting result.

| $^{31}$P NMR | | | | | | | | | | | |
|---|---|---|---|---|---|---|---|---|---|---|---|
| Conditions | | | $R^2$ | Degrees of freedom | Sy.x | $A_0$ (%) | | k ($K^{-1}$) | | $T_{50}$ (K) | |
| DOPE /SOPC | Added lipid | | | | | Value | $CI_{95\%}$ | Value | $CI_{95\%}$ | Value | $CI_{95\%}$ |
| 3:1 | 5% DIM | PC | 0.99 | 18 | 3.6 | 99.4 | ± 3.1 | -0.32 | ± 0.05 | 305.3 | ± 0.5 |
| | | PE | 0.99 | 18 | 2.55 | 84.7 | ± 2.1 | -0.40 | ± 0.05 | 303.5 | ± 0.4 |
| 3:1 | 2.5% DIM | PC | 0.99 | 18 | 4.26 | 96.4 | ± 4.3 | -0.23 | ± 0.04 | 306.5 | ± 0.9 |
| | | PE | 0.99 | 18 | 2.69 | 87,0 | ± 2.7 | -0.26 | ± 0.03 | 304.5 | ± 0.5 |
| 3:1 | 1% DIM | PC | 0.98 | 18 | 4.91 | 99.2 | ± 3.3 | -0.27 | ± 0.06 | 315.0 | ± 0.7 |
| | | PE | 0.98 | 18 | 4.08 | 86.8 | ± 3.8 | -0.19 | ± 0.04 | 311.3 | ± 1.0 |
| 3:1 | 5% DLiPE | PC | 0.98 | 18 | 4.25 | 90.9 | ± 2.3 | -0.51 | ± 0.12 | 317.0 | ± 0.5 |
| | | PE | 0.99 | 18 | 2.91 | 83.4 | ± 1.7 | -0.42 | ± 0.07 | 316.0 | ± 0.4 |
| 3:1 | 5% TAG | PC | 0.99 | 18 | 1.90 | 96.2 | ± 3.2 | -0.10 | ± 0.02 | 321.6 | ± 0.9 |
| | | PE | 0.98 | 18 | 1.65 | 76.3 | ± 1.8 | -0.13 | ± 0.02 | 321.8 | ± 0.8 |
| 3:1 | 2.5% TAG | PC | 0.97 | 18 | 1.91 | 116.7 | ± 14.6 | -0.04 | ± 0.02 | 329.5 | ± 3.5 |
| | | PE | 0.98 | 18 | 1.32 | 94.7 | ± 11.9 | -0.04 | ± 0.02 | 330.6 | ± 3.6 |
| 5:1 | - | PC | 0.99 | 17 | 3.27 | 100.8 | ± 1.2 | -0.33 | ± 0.04 | 314.5 | ± 0.4 |
| | | PE | 0.99 | 17 | 3.09 | 88.3 | ± 2.2 | -0.37 | ± 0.05 | 310.6 | ± 0.5 |
| 5:1 | 5% LysoPC | PC | 0.98 | 17 | 3.63 | 101.0 | ± 2.4 | -0.30 | ± 0.05 | 316.8 | ± 0.5 |
| | | PE | 0.99 | 17 | 2.72 | 93.5 | ± 1.9 | -0.30 | ± 0.04 | 313.8 | ± 0.4 |
| 3:1 | 2.5% DIM + 5% LysoPC | PC | 0.97 | 17 | 4.69 | 93.6 | ± 3.5 | -0.24 | ± 0.05 | 313.6 | ± 0.9 |
| | | PE | 0.98 | 17 | 3.63 | 81.4 | ± 3.5 | -0.18 | ± 0.03 | 311.7 | ± 1.0 |
| 3:1 | 2.5% DIM + 10% LysoPC | PC | 0.99 | 17 | 2.95 | 102.3 | ± 3.7 | -0.11 | ± 0.04 | 329.3 | ± 3.6 |
| | | PE | 0.98 | 17 | 3.15 | 83.7 | ± 3.9 | -0.12 | ± 0.03 | 321.2 | ± 1.5 |
| **CG simulations** | | | | | | | | | | | |
| Conditions | | | $R^2$ | Degrees of freedom | Sy.x | $A_0$ (%) | | k ($K^{-1}$) | | $T_{50}$ (K) | |
| DOPE /SOPC | Added lipid | | | | | Value | $CI_{95\%}$ | Value | $CI_{95\%}$ | Value | $CI_{95\%}$ |
| 3:1 | 5% DIM | PC | 0.99 | 4 | 3.36 | 92.4 | ± 6.59 | -0.57 | ± 1.22 | 308.1 | ± 4.3 |
| | | PE | 0.99 | 4 | 3.31 | 86.3 | ± 6.42 | -0.51 | ± 1.04 | 308.7 | ± 2.9 |
| 3:1 | 2.5% DIM | PC | 0.99 | 5 | 4.91 | 96.3 | ± 6.67 | -0.27 | ± 0.11 | 324.8 | ± 2.2 |
| | | PE | 0.99 | 5 | 5.79 | 93.7 | ± 8.92 | -0.17 | ± 0.08 | 324.1 | ± 3.4 |
| 3:1 | 2.5% DIM + 10% LysoPC | PC | 0.98 | 8 | 6.01 | 88.6 | ± 5.66 | -0.56 | ± 1.04 | 342.5 | ± 4.9 |
| | | PE | 0.97 | 8 | 7.99 | 84.8 | ± 7.56 | -0.49 | ± 1.00 | 342.2 | ± 4.9 |

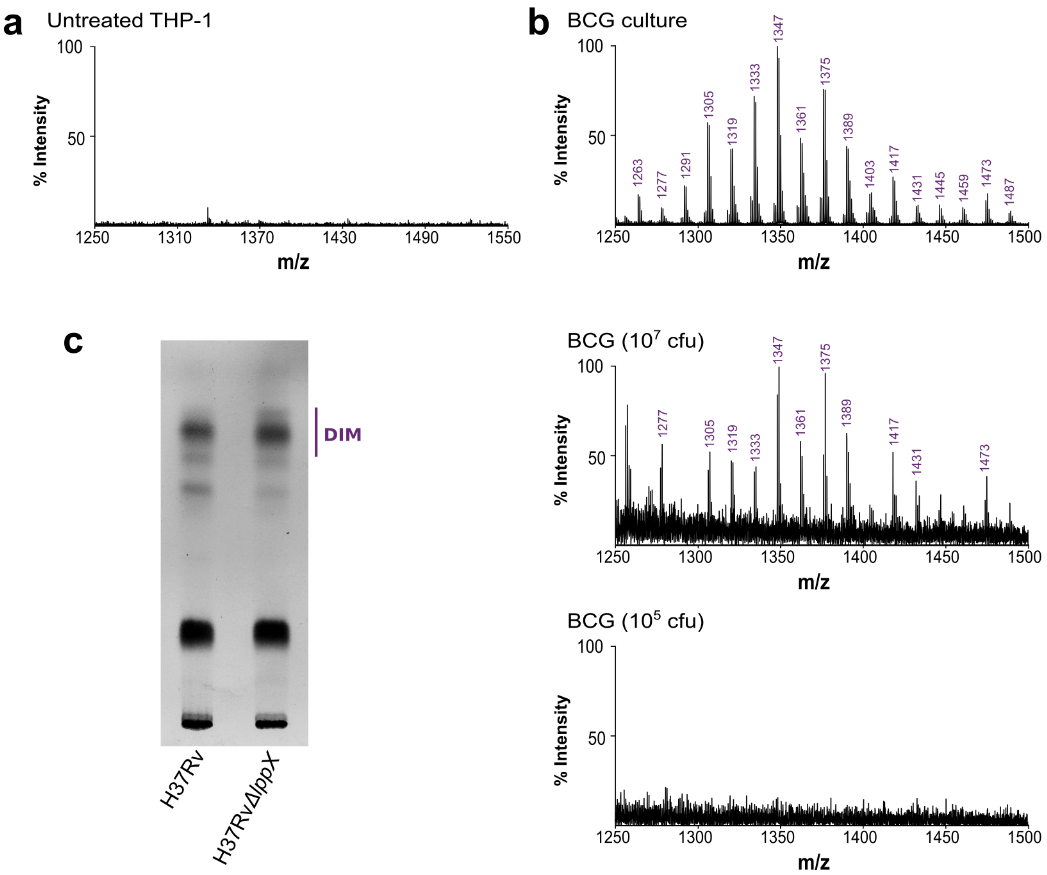


**Fig. S1: DIM lipids are transferred to macrophage membranes.** (**a**) MALDI-TOF spectrum of untreated THP-1 membrane fractions centered on the m/z region where DIM are expected. (**b**) MALDI-TOF spectra of lipid extracts from BCG stock culture and dilutions to $10^7$ resp. $10^5$ cfu. At $10^5$ cfu, the concentration of DIM is below the MALDI-TOF detection threshold. (**c**) Thin layer chromatography analysis (phosphomolybdate staining) of lipids extracted from the WT H37Rv and H37Rv *ΔlppX* strains by the Bligh and Dyer method, showing that the mutant produces a similar amount of DIM as the WT strain.

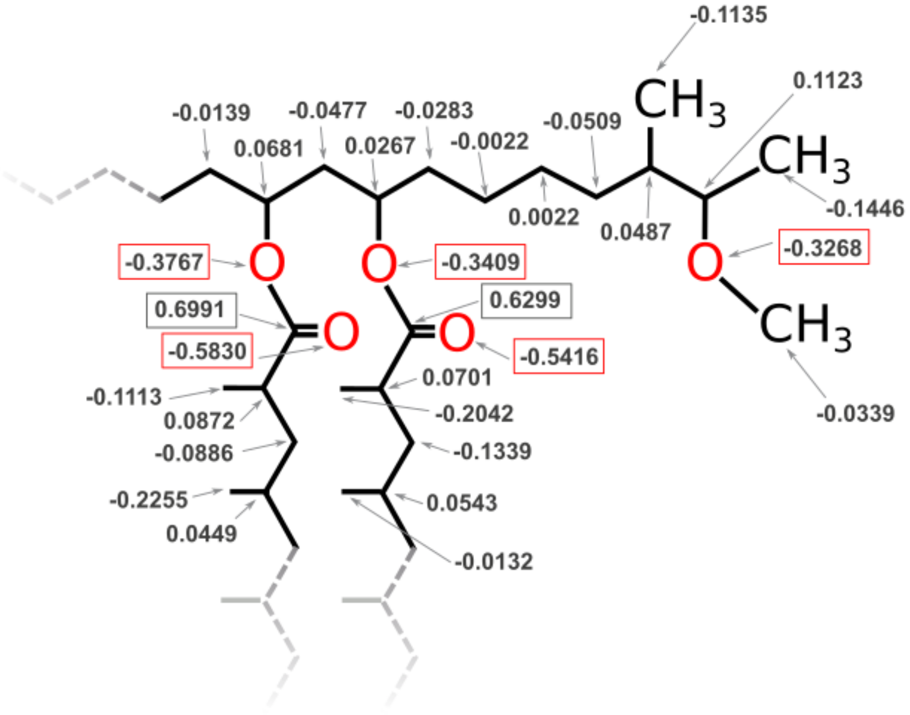


**Fig. S2: DIM partial charges.** Partial charges for the DIM core calculated using the RESP method (see Methods section). For the framed charge values, significant differences were observed compared to the Lipid14 parameters. For the remaining atoms, calculated charges are closed to the ones proposed by standard force fields such as Lipid14.

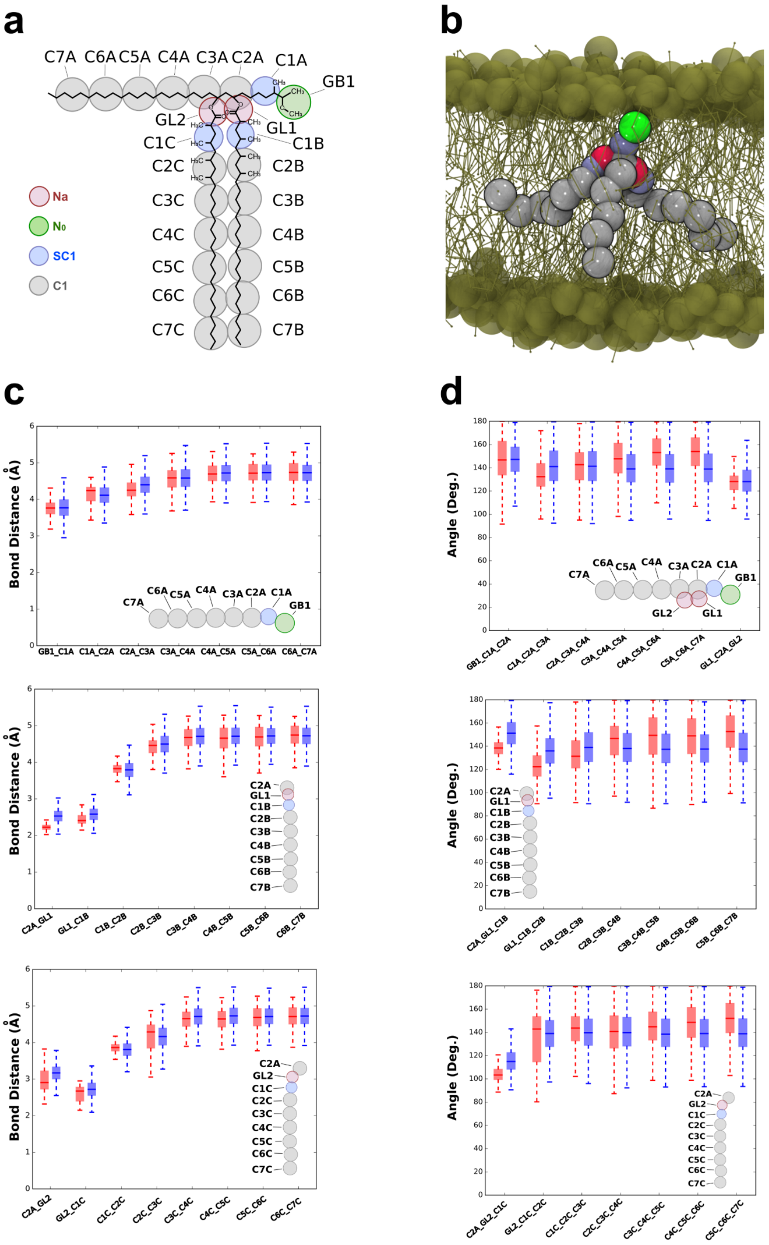


**Fig. S3: DIM coarse grained model parametrization.** (**a**) Mapping between chemical structure and coarse grained model for the DIM molecule. (**b**) DIM molecule in a POPC bilayer. DIM particles are colored as in (a). POPC phosphate particles are displayed in tan spheres. (**c**) Bond length distributions for atomistic (red) and coarse grained (blue) simulations for the phthiocerol moiety and the two mycocerosate tails. (**d**) Angle distributions for atomistic (red) and coarse grained (blue) simulations for the phthiocerol moiety and the two mycocerosate tails.

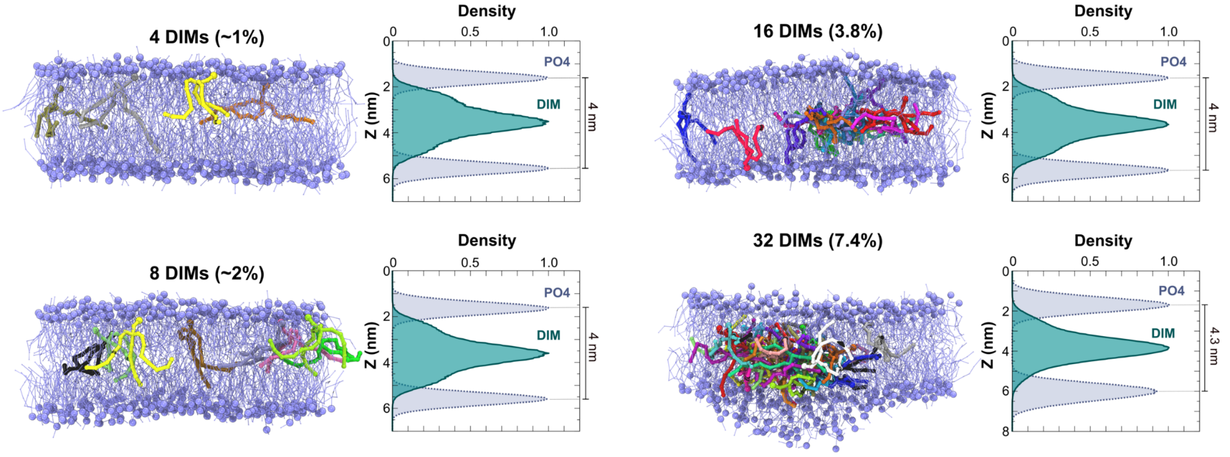


**Fig. S4: DIM aggregation in a POPC bilayer.** Coarse grained model of DIM molecules at different concentrations embedded in a POPC bilayer (see Table S1 for more details). POPC phosphate particles are depicted as blue spheres. On the right, density of DIM lipids and phosphate head groups. Increasing the concentration of DIM molecules drove the DIM aggregation in the inter-leaflet space. At a molar concentration of DIM close to 7%, the DIM aggregation started to increase the POPC membrane thickness.

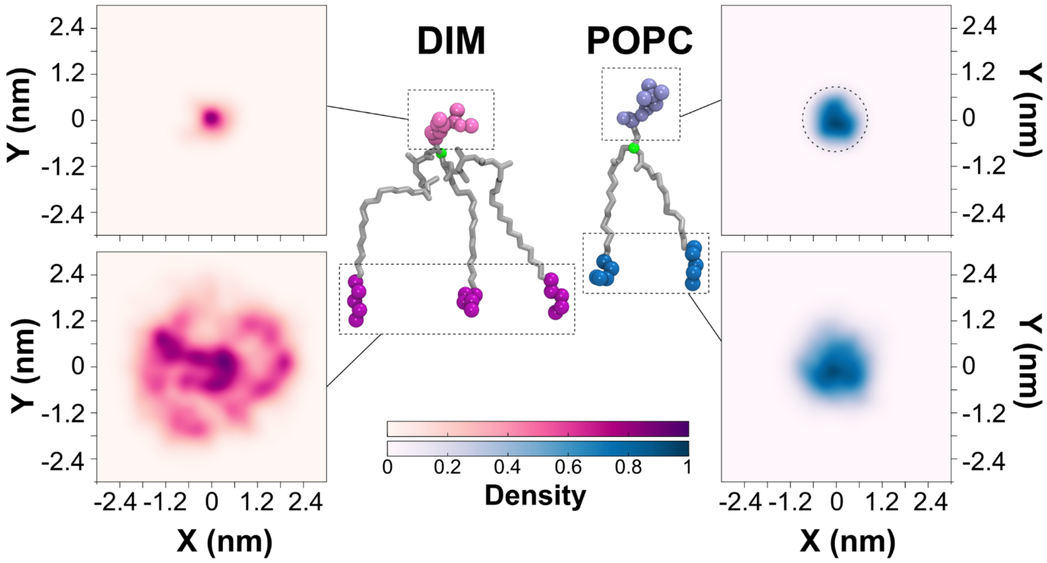


**Fig. S5: Molecular shape of DIM lipids in a POPC bilayer (AT).** 2D density projections of DIM and POPC extremities extracted from atomistic simulations (see **Fig. 2a**). The densities highlight the conical (resp. cylindrical) shape of the DIM (resp. POPC) molecule. Particles depicted in green were used to center the molecules. See also Figure 2 for equivalent results for coarse grained simulations.

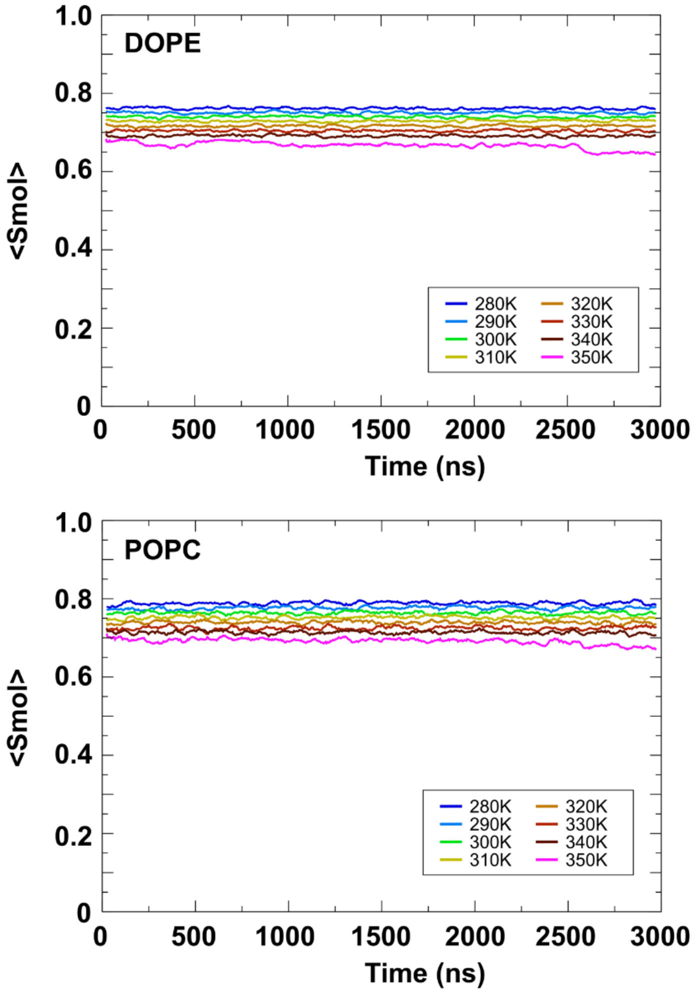


**Fig. S6: 3:1 DOPE-SOPC lipid mixture displays a stable lamellar phase at different temperatures in CG simulations.** Averaged reorientation of the molecular director with respect to the bilayer normal (Smol) for the 3:1 DOPE-SOPC lipid mixture taken during the course of the 3 μs CG simulations (see **Fig. 3b** for an illustration of the system at 320 K). Smol = $< 3\cos^2\theta -1 /2 >$ with $\theta$ the angle between the lipid principal axis and the membrane normal. Smol values for DOPE molecules are slightly lower than the ones for SOPC molecules. This may highlight a higher propensity of DOPE molecules to prefer the non-lamellar phase in comparison of SOPC molecules as seen in NMR experiments (see **Fig. 3c**).

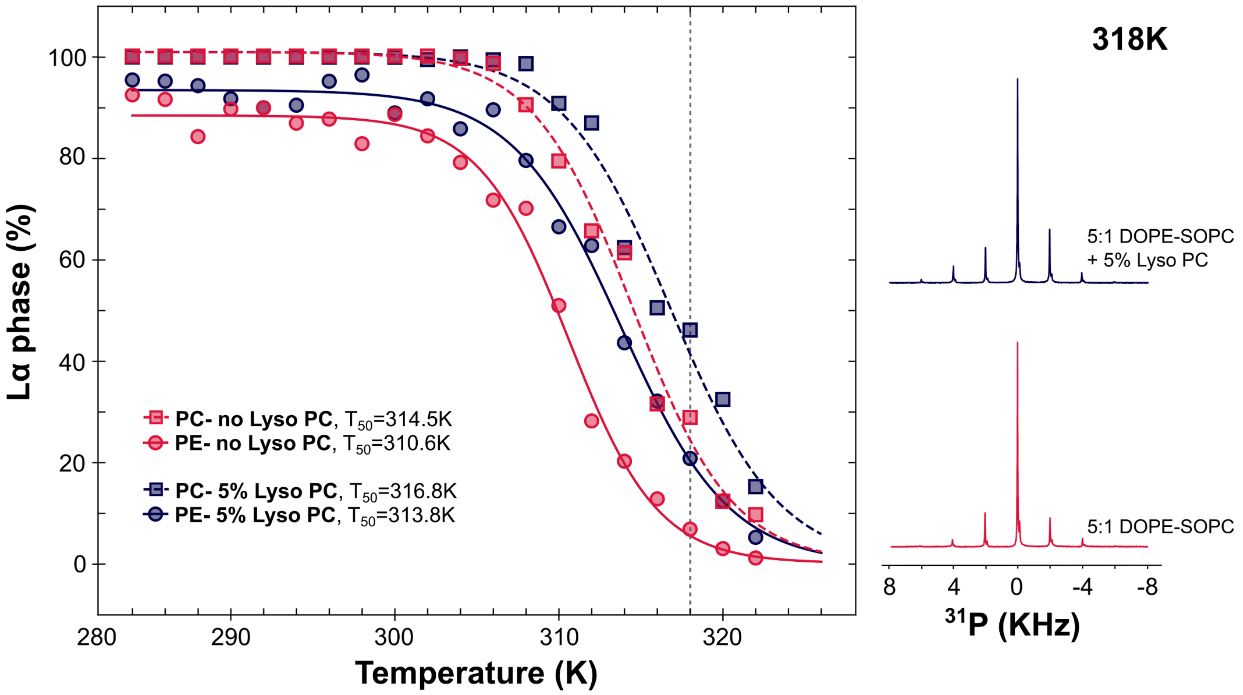


**Fig. S7: LysoPC lipids shift the phase transition of a 5:1 DOPE-SOPC mixture.** Left, evolution of $L\alpha$ to $H_{II}$ phase for the DOPE and SOPC molecules as a function of the temperature with and without addition of 5% of lysoPC lipids. For clarity, error bars were omitted. As seen in **Fig. 3** (see also Methods section), the error was evaluated to +/- 5%. The dashed line highlights the different temperature points obtained for the spectra shown on the right. Right, $^{31}$P NMR spectra for the lipids in a 5:1 DOPE-SOPC lipid mixture with and without addition of 5% of lysoPC at 318 K.

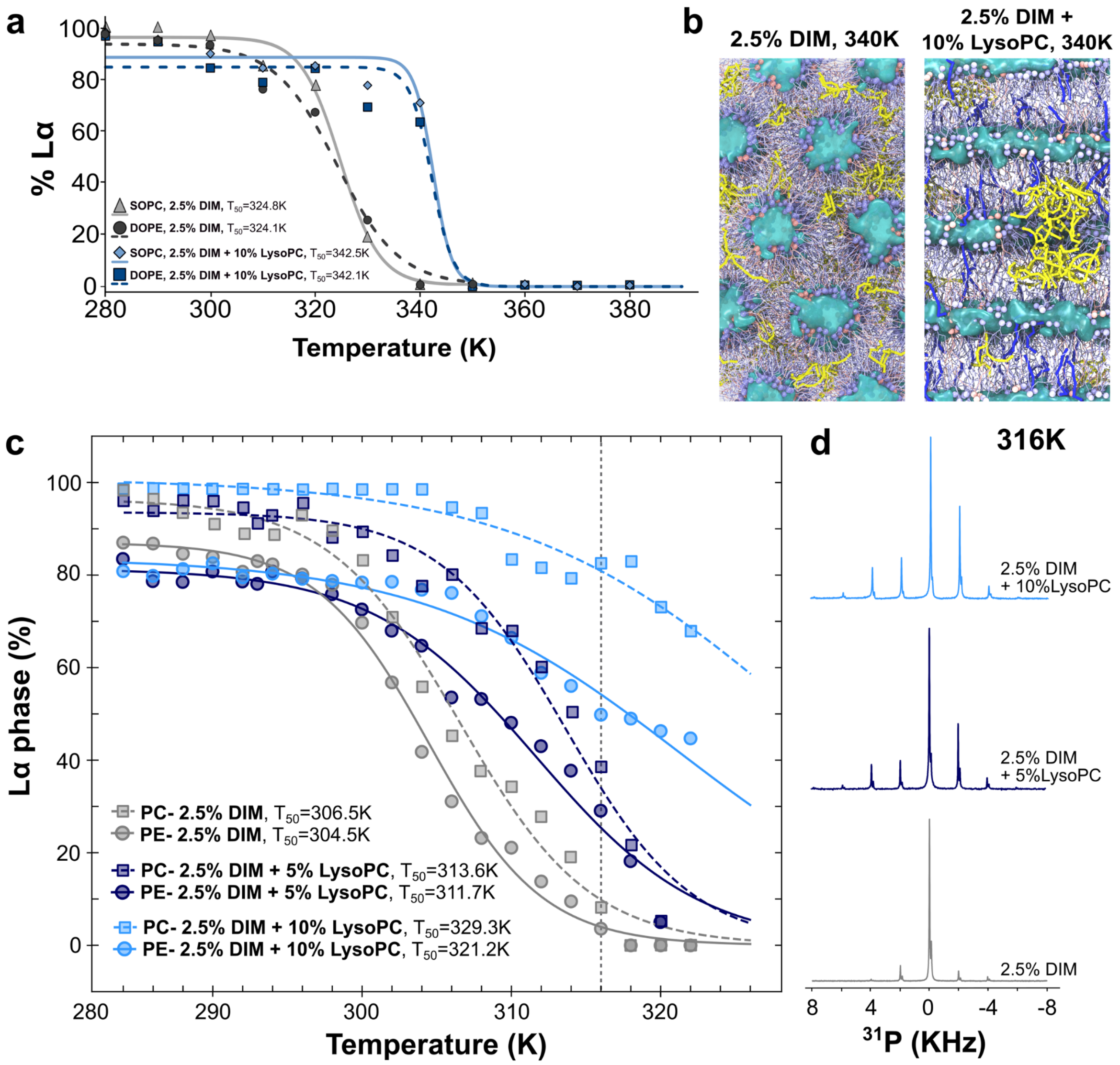


**Fig. S8: LysoPC lipids modulate the curvature action of DIM lipids in a 3:1 DOPE-SOPC mixture.** (**a**) SOPC and DOPE phase transitions calculated from CG-MD simulations with addition of 2.5% of DIM (as presented in **Fig. 3d**) and with addition of 2.5% of DIM and 10% of lysoPC lipids. The addition of 10% of lysoPC lipids shifts the phase transition to higher temperature values highlighting the modulation of the action of DIM by lysoPC molecules. (**b**) Coarse grained models of phase transition for a 3:1 mixture of DOPE-SOPC containing 2.5% of DIM or 2.5% of DIM and 10% of lysoPC. The addition of lysoPC limits the formation and expansion of stalks in simulations. Snapshots taken at the end of the 3µs simulation for a temperature of 340 K. POPC lipids colored in red. DOPE molecules colored in blue. DIM molecules colored in yellow. LysoPC molecules colored in dark blue. Water molecules represented as a blue surface. LysoPC molecules are homogeneously spread throughout the membrane and not concentrated around DIM lipids. (**c**) Evolution of the $L_\alpha$ to $H_{II}$ phase for the DOPE and SOPC molecules as a function of the temperature for a 3:1 DOPE-SOPC mixture containing 2.5% of DIM lipids with different concentrations of lysoPC lipids. For clarity, error bars were omitted. As seen in **Fig. 3** (see also Methods section), the error was evaluated to +/- 5%. The dashed line highlights the different temperature points obtained for the spectra described on the right. Right, $^{31}$P NMR spectra for the lipids in a 3:1 DOPE-SOPC lipid mixture containing 2.5% of DIM lipids with the addition of different concentrations of lysoPC at 316 K.

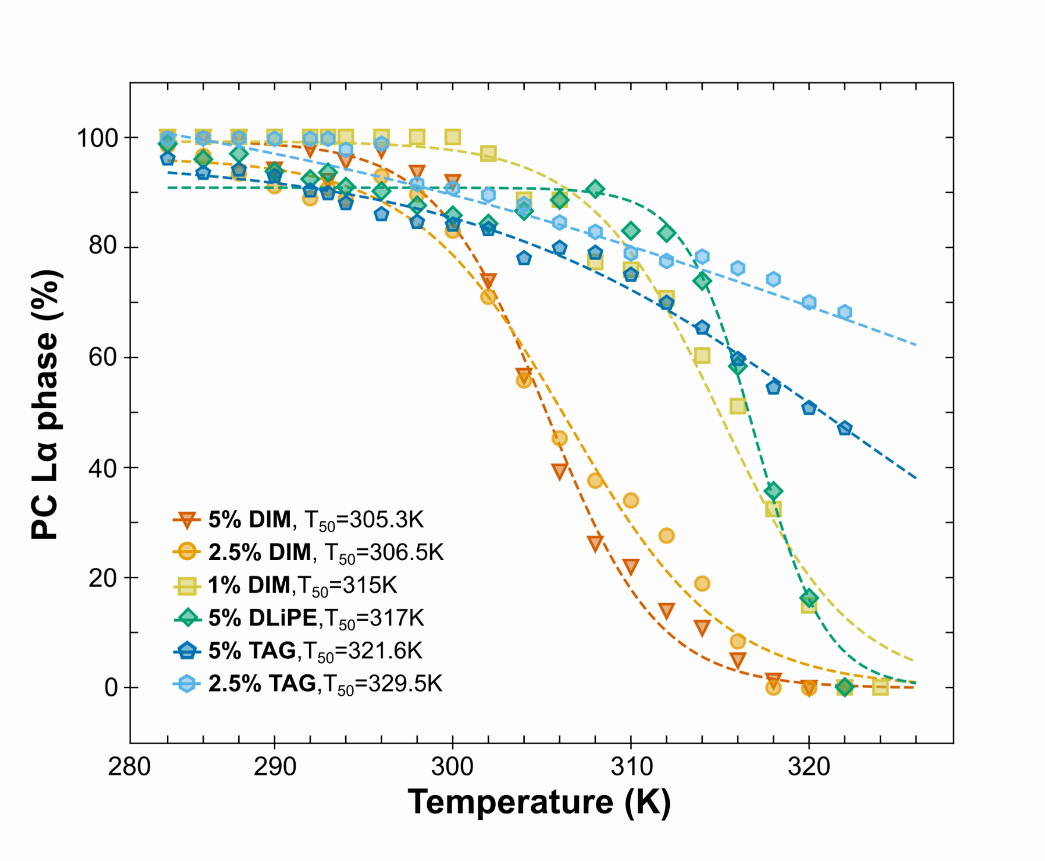


**Fig. S9: Comparison of DIM potency to induce non-bilayer phase with lipids of different shapes ($L_\alpha$ transition of SOPC).** Evolution of the percentage of $L_\alpha$ phase for the SOPC molecules as a function of temperature upon incorporation of different concentrations of DIM, DLiPE and TAG in a (3:1) DOPE/SOPC mixture (see also **Fig. 4** for the corresponding curves for DOPE molecules). For clarity, error bars were omitted. As seen in **Fig. 3** (see also Methods), the error was evaluated to +/- 5%.

**Supplementary Movie S1:** CG-MD simulation of the $L_\alpha$-to-$H_{II}$ phase transition of a DOPE/SOPC (3:1) mixture induced by DIM lipids (red). Water molecules are depicted in transparent blue. For DOPE and SOPC lipids, the polar head group particle PO4 is depicted in green while ester particles (GL1 and GL2) are depicted in gray. For clarity reasons, the SOPC and DOPE acyl chains were removed. A high resolution version is available at: http://matthieuchavent.com/videos/movie_S1.mov
